# Dopaminergic inhibition of the inwardly rectifying potassium current in direct pathway medium spiny neurons in normal and parkinsonian striatum

**DOI:** 10.1101/2024.04.29.590632

**Authors:** Qian Wang, Yuhan Wang, Francesca-Fang Liao, Fu-Ming Zhou

**Affiliations:** Department of Pharmacology, Addiction Science and Toxicology, College of Medicine, University of Tennessee Health Science Center, Memphis, TN 38103, USA

**Keywords:** basal ganglia, dopamine receptor, inwardly rectifying potassium channel, medium spiny neuron, Parkinson’s disease, striatum

## Abstract

Despite the profound behavioral effects of the striatal dopamine (DA) activity and the inwardly rectifying potassium channel (**Kir**) being a key determinant of striatal medium spiny neuron (MSN) activity that also profoundly affects behavior, previously reported DA regulations of Kir are conflicting and incompatible with MSN function in behavior. Here we show that in normal mice with an intact striatal DA system, the predominant effect of DA activation of D1Rs in D1-MSNs is to cause a modest depolarization and increase in input resistance by inhibiting Kir, thus moderately increasing the spike outputs from behavior-promoting D1-MSNs. In parkinsonian (DA-depleted) striatum, DA increases D1-MSN intrinsic excitability more strongly than in normal striatum, consequently strongly increasing D1-MSN spike firing that is behavior-promoting; this DA excitation of D1-MSNs is stronger when the DA depletion is more severe. The DA inhibition of Kir is occluded by the Kir blocker barium chloride (BaCl_2_). In behaving parkinsonian mice, BaCl_2_ microinjection into the dorsal striatum stimulates movement but occludes the motor stimulation of D1R agonism. Taken together, our results resolve the long-standing question about what D1R agonism does to D1-MSN excitability in normal and parkinsonian striatum and strongly indicate that D1R inhibition of Kir is a key ion channel mechanism that mediates D1R agonistic behavioral stimulation in normal and parkinsonian animals.

## 1. Introduction

Dopaminergic (DA) activity is highly concentrated in the striatum with an intense DA innervation and a high level of expression of DA D1 and D2 receptors (Gerfen and Bolam 2017). This DA activity is absolutely required for normal motor function: Loss of the striatal DA or blockade of striatal DA receptors leads to loss of motor function in both humans and animals; DA and D1R- like (D1R hereafter) agonists profoundly stimulate motor activity (LeWitt and Fahn 2016; Li and Zhou 2013, Mailman et al. 2001, Rascol et al. 1999). However, the ion channel and cellular mechanisms underlying DA’s spectacular motor stimulation are poorly understood with conflicting results in the literature (Nicola et al. 2000; Humphries and Prescott 2010). More recent studies have shown that D1R-bypassing optogenetic activation of the D1R-expressing medium spiny neurons (D1-MSNs) promotes motor function (Friend and Kravitz 2014; Sippy et al. 2015), but critical and long-standing questions remain: How does D1R agonism (the endogenous mechanisms) affect D1-MSN intrinsic excitability that can affect the spiking activity of these neurons, and what ion channels are critically involved, and what is the key ion channel that mediates DA-D1R agonistic motor stimulation? Delineating these fundamental mechanisms will advance our understanding of basal ganglia physiology and PD pathophysiology and treatment.

Since the inwardly rectifying potassium (Kir; molecularly Kir2 type, mainly Kir2.3 subtype) channel is highly expressed and tonically active in MSNs and a key determinant of MSN intrinsic excitability (Nisenbaum and Wilson 1995; Shen et al. 2007), Kir is a key candidate for DA regulation of MSN activity. Multiple studies have investigated how D1R agonism affects MSN intrinsic excitability and reported that D1R agonism caused hyperpolarization and increased the Kir-mediated inward rectification and hence decreased intrinsic excitability and spike firing in MSNs in dorsal striatum (DS) and nucleus accumbens (NAc) (Higashi et al. 1989; Nicola et al. 2000; Pacheco-Cano et al. 1996; Uchimura et al. 1986). But these results were confounded because recordings were made in mixed, unidentified D1- and D2-MSNs, D1Rs are expressed only in 50% of the MSNs and thus it is impossible to have a true D1R response in all recorded MSNs as reported (e.g. Pacheco-Cano et al. 1996). A more recent study in identified D1-MSN and D2-MSNs reported that D1R agonism *<u>enhanced</u>* Kir and decreased D1-MSN excitability and D2R agonism did the opposite in D2-MSNs (Zhao et al. 2016). However, these reported DA effects are incompatible with the function of these MSNs in animal behavior: optogenetic activation of D1-MSNs stimulates motor behavior (Friend and Kravitz 2014; Kravitz et al. 2010; Sippy et al. 2015) and striatal D1 agonist microinjection stimulates movement (Wang and Zhou 2017). A more recent study (Lahiri and Bevan 2020) also did not resolve the question, as discussed in the Discussion section.

Additionally, previous studies have reported contradictory effects of DA loss on intrinsic membrane properties such as resting membrane potential (RMP) and input resistance (R_In_) measured in the same cell type in the same animal model. For example, Lieberman et al. (2018) reported that D1-MSNs had more depolarized RMP and higher R_In_ in Pitx3Null mice than in wild-type (WT) mice, but Suarez et al. (2018) reported that D1-MSNs had similar RMP and R_In_ in Pitx3Null mice and WT mice. It is no wonder that Humphries and Prescott (2010) lamented that the reported DA effects on striatal MSNs have been “in a state of permanent controversy.” Thus, new studies are needed to determine the true DA effects in D1-MSNs and the key underlying ion channel mechanism.

## 2. Materials and methods

### 2.1. Animals

#### Transcription factor Pitx3-/- null mutant (Pitx3Null) mice

We have used this spontaneous mutant mouse line extensively and have a deep understanding about this mouse (Ding et al. 2015a; Li et al. 2013; Li et al. 2015; Li and Zhou 2013; Sagot et al. 2018; Wei et al. 2013, 2017; Wang and Zhou 2017). Pitx3 is required for the survival of most DA neurons in the substantia nigra, whereas about 50% DA neurons in the ventral tegmental area do not require Pitx3 to survive; hence these mice have a severe and selective DA neuron loss in the substantia nigra and a severe DA denervation in the dorsal striatum, resembling the DA loss pattern in the PD brain (**Fig. 1A,B**) (Ding et al. 2015a; Li et al. 2013; Li et al. 2015; Li and Zhou 2013; Sagot et al. 2018; Wei et al. 2013, 2017; Wang and Zhou 2017; Zhong et al. 2023). In our prior studies, we observed no sex difference in DA denervation and response to L-dopa in Pitx3Null mice.

**Fig. 1.**
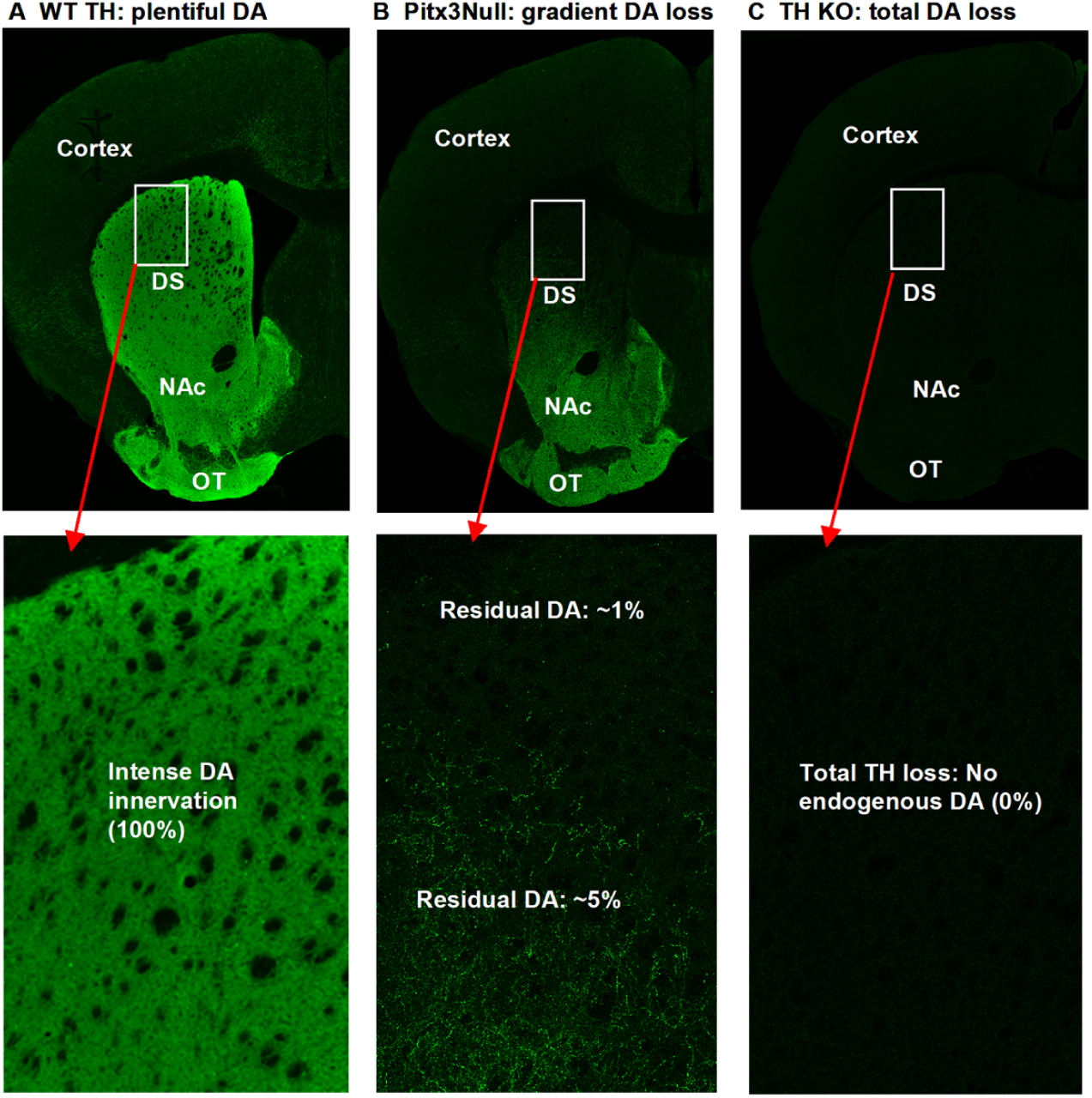
Different DA denervation patterns of Pitx3Null mice and TH KO mice. **A.** Normal WT mice have an intense DA innervation in the entire striatum, as indicated by the intense TH immunostaining signal. **B.** Pitx3Null mice have a dorsal-ventral DA loss gradient in the striatum. **C.** TH KO mice have a total lack of TH in the entire striatum. The boxed areas are expanded and displayed below the main images.

Two breeding pairs of heterozygous Pitx3+/- mice (stock # 000942) were purchased from the Jackson Laboratory (Bar Harbor, ME), resulting in a small colony of homozygous Pitx3-/- (Pitx3Null), heterozygous Pitx3+/-, and wild-type Pitx3+/+ (Pitx3WT) mice at the animal facility at UTHSC in Memphis. Pitx3Null mice are aphakic and thus clearly identified (Wei et al. 2013; Zhou et al. 2016). The genotypes were also determined by PCR-based genotyping to identify WT, homozygotes, and heterozygotes (Li et al. 2013).

#### DA neuron-selective tyrosine hydroxylase (TH) KO mice with hyperfunctional DA receptors

In these mice, the **TH** gene in DA neurons is selectively deleted (Zhou and Palmiter 1995). Thus, there is a lack of TH and DA in the entire striatum (**Fig. 1A,C**); but the DA axons are intact and can make DA from L-dopa. An established feature of these mice is their hyperfunctional D1Rs and D2Rs (Kim et al. 2000, Seeman et al. 2005; Zhong et al. 2023).

Breeders for these mice was generously donated by Dr. Martin Darvas, University of Washington (Wang et al. 2020). This mouse is now also available at the Jackson Laboratory: Stock # 009688 [B6;129-Dbh^tm2(Th)Rpa^ Th^tm1Rpa^/J]. TH KO mice are identified by PCR-based genotyping and by their baseline akinesia and their robust motor response to IP injected L-dopa.

Additionally, two pairs of heterozygous BAC D2-GFP+/- breeder mice (Gong et al., 2003) were purchased from the Mutant Mouse Research Resource Centers (MMRRC) ([stock Tg(Drd2-EGFP) S118Gsat/Mmnc (000230-UNC); RRID: MMRRC_000230-UNC]) and crossed to TH KO mice and Pitx3Null mice, eventually producing TH KO-D2-GFP mice and Pitx3Null- D2-GFP mice, enabling recording identified MSNs in brain slices. Mice had free access to food and water. All procedures were approved by The Institutional Animal Care and Use Committee of The University of Tennessee Health Science Center (animal protocol # 17-033.0).

### 2.2. Patch clamp electrophysiology

#### Brain slice preparation

Coronal brain slices containing the striatum were prepared using a conventional method described before (Wei et al. 2013; Wei et al. 2017). Mice of postnatal (PN) days 18-21 were used for the majority of our experiments; PN28-30 mice were also used to verify that conclusions from PN20 mice are still correct at a more mature age. These mice were euthanized by decapitation, and their brains were dissected out quickly and immediately immersed in an oxygenated ice-cold cutting solution (containing, in mM, 220 glycerol, 2.5 KCl, 1.25 NaH_2_PO4, 25 NaHCO_3_, 0.5 CaCl_2_, 7 MgCl_2_, 20 D-glucose) for 2 min. Three hundred-micrometer-thick brain slices were cut on a Leica Zero Z VT1200S vibratome (Leica Microsystems, Wetzlar, Germany). The slices were transferred to a holding chamber filled with a standard extracellular bathing solution (containing, in mM, 125 NaCl, 2.5 KCl, 25 NaHCO_3_, 1.25 NaH_2_PO_4_, 2.5 CaCl_2_, 1.3 MgCl_2_, and 10 D-glucose) that was continuously bubbled with 95% O_2_- 5% CO_2_ at 34°C for 30 min (this solution was also used for current clamp recording), and then the holding chamber was kept at room temperature (25°C). Brain slices were used within 1–6 hours after being prepared. Ascorbic acid (vitamin C, 0.4 mM) was added to all solutions to protect the brain tissue.

#### Brain slice-patch clamp recording

Brain slices were placed in a recording chamber and continuously perfused at 2 ml/min with the standard extracellular bathing solution saturated with 95% O_2_-5% CO_2_. Recordings were made under visual guidance of a video microscope (Olympus BX51WI and Zeiss Axiocam MRm digital camera) equipped with Nomarski optics and a ×60 water immersion lens. D1-MSNs were identified as D2-GFP negative cells in transgenic D2-eGFP mice. This identification is reliable based on the fact that D1Rs and D2Rs are overwhelmingly if not completely segregated, especially in DS and NAc, the striatal areas this study targeted (Valjent et al. 2009). Our prior study further indicated that D1-MSNs so identified have a D1R-mediated presynaptic facilitation whereas D2-MSNs have a D2R-mediated presynaptic inhibition (Wei et al. 2017). Cholinergic interneurons are characteristically large neurons and easily avoided; GABAergic interneurons are significantly larger than MSNs, can be recognized by trained eyes, and have membrane properties distinct from those of MSNs (Kawaguchi 1993; Koos and Tepper 1999; Nisenbaum and Wilson 1995; Tepper et al. 2004; Wei et al. 2017). Thus, our identification of D1-MSNs was reliable.

A Multiclamp 700B amplifier, pClamp 9.2 software, and Digidata 1322A interface (Molecular Devices, Sunnyvale, CA) were used to acquire data. Patch pipettes were pulled from borosilicate glass capillary tubing (1.1-mm ID, 1.5-mm OD, purchased from Sutter instrument, Novato, CA: cat. # BF150-110-10) using a PC-10 puller (Narishige, Tokyo, Japan) and had resistances of 4 MΩ for recording. All recordings were made at 30–32°C.

The intracellular solution used for recording membrane potential and currents contained (in mM) 135 KCl, 0.5 EGTA, 10 HEPES, 2 Mg-ATP, 0.2 Na-GTP, and 4 Na_2_-phosphocreatine with pH 7.25 and 285 mOsm. To voltage clamp-record Kir and other K currents, we used the following relatively high KCl and zero Ca^2+^ extracellular solution containing (in mM) 125 NaCl, 3.5 KCl, 25 NaHCO_3_, 1.25 NaH_2_PO4, 3.8 MgCl_2_, and 10 D-glucose. The calculated K reversal potential at –87.5 mV for our extracellular and intracellular solution pair. Tetrodotoxin (TTX; 1 μM) was added to the extracellular solution for recording K currents. All recordings were made in the presence of 6,7-dinitro-quinoxaline-2,3-dione (DNQX; 10 μM), d,l-2-amino-5- phosphonovalerate (AP5; 20 μM), and 100 μM picrotoxin in extracellular bathing solution to block ionotropic glutamate receptors and GABA_A_ receptors. Our pilot experiment indicated that D2 antagonist eticlopride did not affect the DA excitation of D1-MSNs in TH KO mice, confirming that the DA effects recorded in D1-MSNs were direct effects on D1-MSNs.

Electrical signals were filtered at 10 kHz using the built-in four-pole low-pass Bessel filter in the Multiclamp 700B patch-clamp amplifier and digitized at 20 kHz using Digidata 1322A. At least five mice were used to obtain an averaged electrophysiological data point, with each mouse yielding one or two useful cells. The Clampfit 9.2 software was used to analyze current clamp and voltage clamp data for membrane potentials, input resistance, action potentials and Kir.

### Extraction of Kir

The voltage-dependence or the I–V relationship of the DA-sensitive current or Kir was determined by using a linear voltage ramp ranging from –120 mV to –40 mV at a speed of 0.15 mV/ms, followed by a digital subtraction procedure (DA-sensitive K current = control current – current under DA). Because Kv currents are large and cannot be selectively blocked without inhibiting Kir and because the recording conditions before and during DA application usually cannot be absolutely identical, especially access resistance, such that the subtraction-extracted difference Kir current (under voltage clamp) and its I–V become unreliable when the difference Kir is small, e.g., only 20 pA. But current clamp recording is less sensitive to access resistance change. Thus, we focused on current clamp experiments in normal WT animals. Certainly this technical problem became small when the DA effects on Kir were large in parkinsonian striatum, allowing us to perform both current clamp and voltage clamp experiments to delineate DA-induced changes in Kir in D1-MSNs.

### 2.3. Unilateral intrastriatal microinjection and contralateral rotation

For this set of experiments, we used 4-5 month old TH KO mice. Bilateral 26-gauge microinjection guide cannulas were implanted with the guide cannula tip being 1 mm above the target area in both sides of the dorsal striatum, as we have used in our prior studies (Wang and Zhou 2017; Wang et al. 2020). An individual internal cannula that is 1 mm longer than implanted guide cannulas was used to reach our target brain area, the dorsal striatum with the coordinates being AP 0 mm, ML ± 2 mm from bregma and DV -3.0 mm from the surface of skull. After a recovery of at least 10 days, behavioral testing was always performed in the afternoon. On the testing day, a tiny dose of L-dopa (1 mg/kg) with benserazide was given to the TH KO mouse by intraperitoneal injection 30min before behavioral testing such that the mouse had the physical strength to rotate when the activity and function of the two side of the striatum became unbalanced upon unilateral drug microinjection into the striatum. The mouse was placed in a 24 cm × 24 cm square recording cage and was video-recorded, starting at 10 min before drug injection and ending at 60 min after unilateral drug microinjection. In this study, 0.2 μg BaCl2- 2H2O (i.e. 0.2 μL 4 mM solution), 0.2 μg SKF81297, and a mixture of 0.2 μg BaCl2 and 0.2 μg SKF81297 (all in the fixed 0.2 μL saline) were slowly microinjected into a single side of the dorsal striatum at the speed of 40 nL/min, inducing unilateral rotations. As a motor behavioral readout, rotations can be easily recognized and was manually counted and binned into 10-min segments. A mouse received only one microinjection on the testing day, and an interval of at least 2 days was given to allow the mouse to recover fully, before the next microinjection.

### 2.4. Immunohistochemistry

Conventional immunofluorescence methods were used to detect DA axons in the striatum (Zhou et al. 2009; Li and Zhou 2013; Wei et al. 2013; Ding et al. 2015). The brains were fixed in 4% paraformaldehyde dissolved in phosphate buffered saline (PBS) at 4°C overnight and then sectioned on a vibratome. The free-floating sections (50 μm in thickness) were incubated with 2% fat-free milk, 1% bovine serum albumin, and 0.4% Triton X-100 in PBS for 1 h at room temperature to block nonspecific binding and permeabilize the cell membrane, respectively. After thorough rinsing, the free-floating sections were incubated for 48 h at 4°C with the primary antibody, a polyclonal tyrosine hydroxylase antibody raised in rabbit (diluted at 1:1000; Novus Biologicals, Littleton, CO; cat. # NB300-109), and then rinsed in PBS, followed by incubating with a donkey anti-rabbit secondary antibody conjugated with the green Alexa Fluor 488 (diluted at 1:400; Invitrogen), for 3 h at room temperature. For D1R staining, the primary D1R antibody is a rat monoclonal antibody purchased from Sigma (cat. # D-187, Sigma, St. Louis, MO; used at 1:500 dilution). Donkey anti-rabbit secondary antibody conjugated with green Alexa Flour 488 and donkey anti-rat secondary antibody conjugated with green Alexa Flour 488 were purchased from Invitrogen (cat. # A21206 and cat. # A21208). Confocal microscopic images were acquired using a Zeiss 710 laser scanning confocal microscope and the associated image acquisition and processing software Zen.

### 2.5. Drugs and chemicals

6,7-dinitroquinoxaline-2,3-dione (DNQX) (cat. # 0189), 2-amino-5-phosphonopentanoic acid (AP5) (cat. # 0106) and picrotoxin (cat. # 1128) were purchased from Tocris. Barium chloride (BaCl2-2H2O) (cat. # 217565) was purchased from Sigma-Aldrich (St. Louis, MO). Dopamine HCl was purchased from Sigma-Aldrich. Forskolin was purchased from MedchemExpress.com. Tetrodotoxin was purchased HelloBio.com (#HB1035, Princeton, NJ). Other routine chemicals were purchased from Sigma-Aldrich (St. Louis, MO). SKF81297-HBr were purchased from Tocris (cat. # 1447).

We need to note here that since D1Rs and D2Rs are segregated in the 2 groups of MSNs (Valjent et al. 2013), bath application of DA will activate only D1Rs in D1-MSNs and separately D2Rs in D2-MSNs in a coronal brain slice preparation with NMDA and non-NMDA glutamatergic receptors and GABA_A_ receptors blocked, but D2R activation does not affect D1- MSNs under our recording condition. We chose DA because it is the endogenous neurotransmitter for DA receptors and hence more functionally relevant than selective exogenous DA agonists.

### 2.6. Statistical analysis

Values were expressed as mean ± SEM. The paired Student’s t test was used to compare changes in the same groups of cells before and during drug application. The un-paired t-test was used to compare the measurements of the same parameter before and after drug treatment in the same mouse group. One-way ANOVA and post hoc Tukey test were used to compare drug-induced rotations 10 min after drug injection in TH KO mice. Following the recent professional guidance on statistics (Amrhein et al. 2019; Wasserstein et al. 2019), p values are given in the text, the figures, or the table, but the traditional binary threshold at p = 0.05 and the phrases “statistically significant” and “statistically not significant” are not used; instead, we base our conclusions on the totality of the data.

## 3. Results

### 3.1. DA modestly increases D1-MSN excitability in the <u>dorsal striatum (DS)</u> in normal mice with intact DA innervation

We first examined DA effects on D1-MSN intrinsic excitability and spike firing, in brain slices with fast glutamatergic and GABA receptors blocked, in DA innervation-intact DS and NAc in normal mice. To increase recording quality and data reliability, we started with PN18-21 mice that yield more viable brain slices and hence high quality recordings. After obtaining a stable baseline recording in current clamp mode, 10 μM DA was bath-applied. We observed that 10 μM DA had modest but consistent excitatory effects on these D1-MSNs by causing a depolarization, increasing the whole-cell input resistance (R_In_), and increasing the membrane charging time constant (τ), and increasing the number of depolarizing current pulse-triggered spikes. Examples and pooled data ― scatter plots of these effects are displayed in **Fig. 2A** and the numerical values are listed in **Table 1**. Bath application of 10 μM DA did not affect the threshold membrane potential for action potential firing, the action potential waveform, or the afterhyperpolarization. Similar DA effects were observed in D1-MSNs in DS in PN28-30 WT mice (**Fig. 2B**). As evident in **Fig. 2A1-A3** and **Fig. 2B1-B3**, a consistent DA effect was a reduction in the inward rectification and this effect became larger as the membrane potential became more negative, indicating a mediation of DA-triggered inhibition of Kir, although we were unable to obtain reliable voltage clamp I-V data because this potential DA agonism- inhibited Kir was likely to be <20 pA (more details in the method section: **Extraction of Kir)**.

**Fig. 2.**
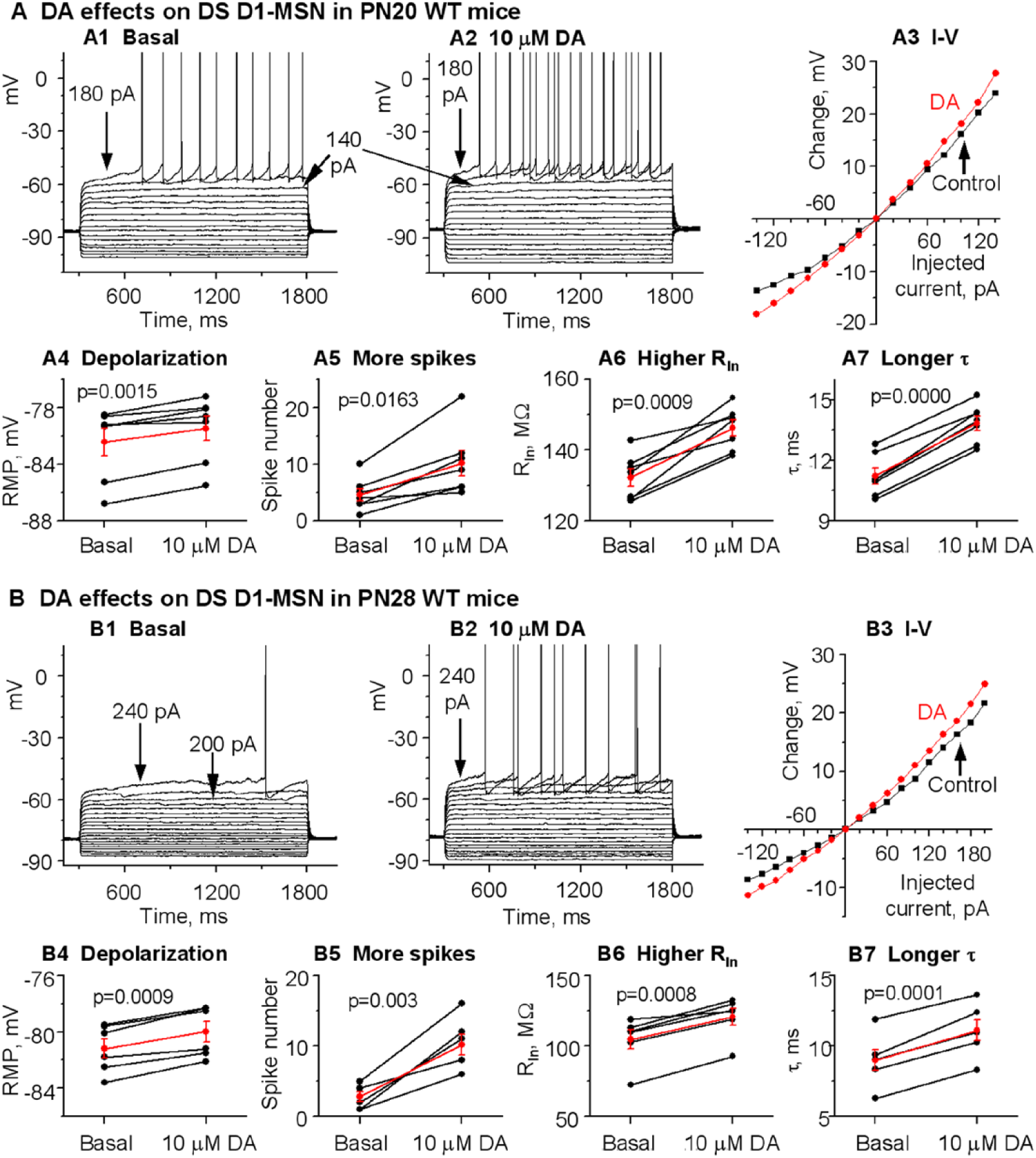
DA normally and modestly increases D1-MSN excitability in the **dorsal striatum (DS)** in PN18-20 and PN28-30 **WT mice** with intact DA innervation. **A.** An example PN20 D1-MSN receiving hyperpolarizing and depolarizing current injection before (**A1**) and during (**A2**) bath application of 10 μM DA. Current pulse-induced voltage changes are quantified in **A3**. **A4-A7** are pooled data showing the DA effect on RMP (**A4**), action potential firing (**A5**), RIn (**A6**) and τ (**A7**) in 7 D1-MSNs. p values are from paired t-tests. **B.** An example PN28 D1-MSN receiving hyperpolarizing and depolarizing current injection before (**B1**) and during (**B2**) bath application of 10 μM DA. Current pulse-induced voltage changes are quantified in **B3**. **B4-B7** are pooled data showing the DA effect on RMP (**B4**), action potential firing (**B5**), RIn (**B6**) and τ (**B7**) in 6 D1-MSNs. p values are from paired ttests.

**Table 1.**
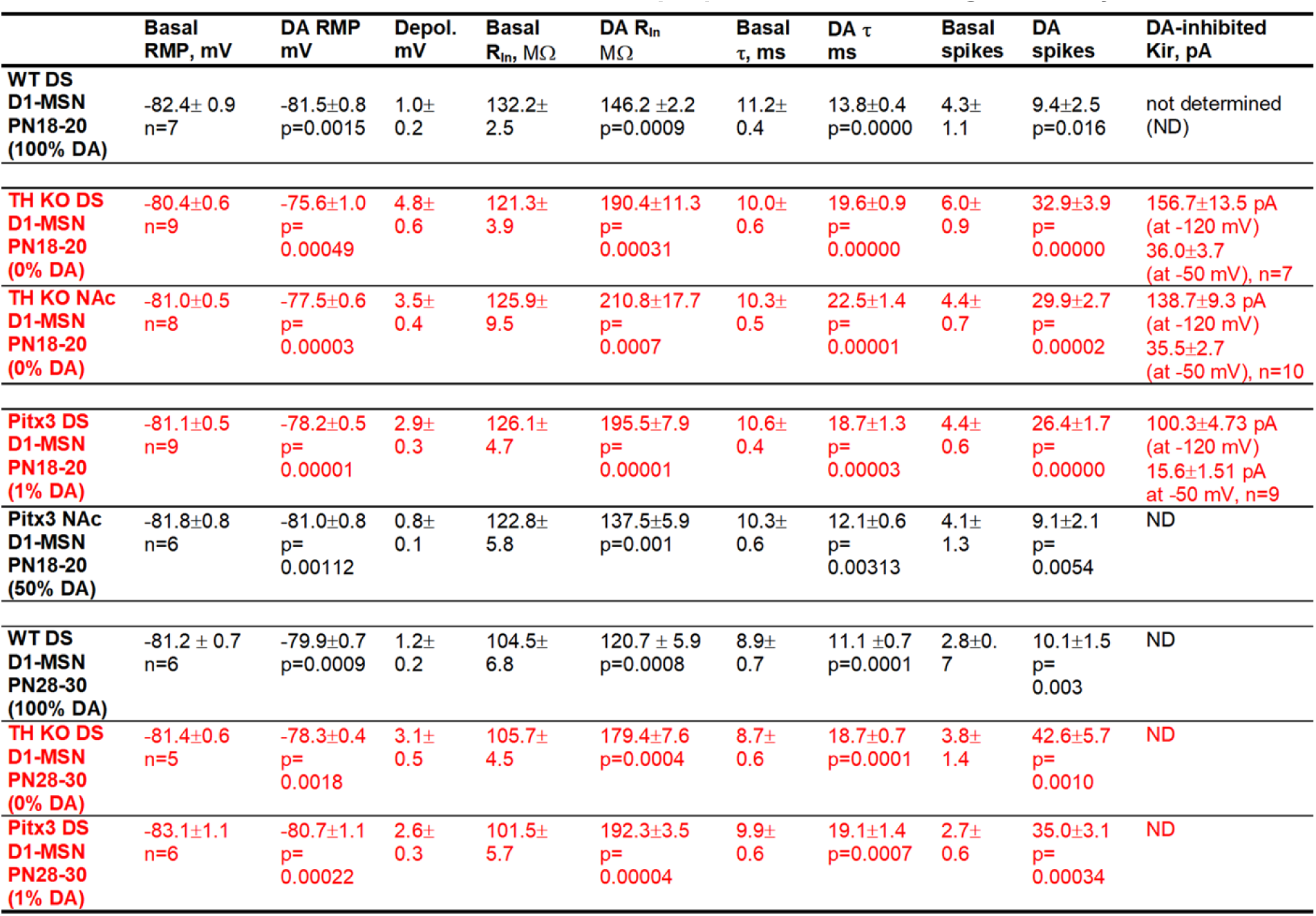
D1-MSN intrinsic membrane properties and their regulation by DA.

|  |  |  |  |  |  |  |  |  |  |  |
| --- | --- | --- | --- | --- | --- | --- | --- | --- | --- | --- |
| | Basal RMP, mV | DA RMP mV | Depol. mV | Basal $R_{in}$, MΩ | DA $R_{in}$ MΩ | Basal $\tau$, ms | DA $\tau$ ms | Basal spikes | DA spikes | DA-inhibited Kir, pA |
| **WT DS**<br>**D1-MSN**<br>**PN18-20**<br>**(100% DA)** | -82.4± 0.9<br>n=7 | -81.5±0.8<br>p=0.0015 | 1.0±<br>0.2 | 132.2±<br>2.5 | 146.2 ±2.2<br>p=0.0009 | 11.2±<br>0.4 | 13.8±0.4<br>p=0.0000 | 4.3±<br>1.1 | 9.4±2.5<br>p=0.016 | not determined<br>(ND) |
| **TH KO DS**<br>**D1-MSN**<br>**PN18-20**<br>**(0% DA)** | -80.4±0.6<br>n=9 | -75.6±1.0<br>p=<br>0.00049 | 4.8±<br>0.6 | 121.3±<br>3.9 | 190.4±11.3<br>p=<br>0.00031 | 10.0±<br>0.6 | 19.6±0.9<br>p=<br>0.00000 | 6.0±<br>0.9 | 32.9±3.9<br>p=<br>0.00000 | 156.7±13.5 pA<br>(at -120 mV)<br>36.0±3.7<br>(at -50 mV), n=7 |
| **TH KO NAc**<br>**D1-MSN**<br>**PN18-20**<br>**(0% DA)** | -81.0±0.5<br>n=8 | -77.5±0.6<br>p=<br>0.00003 | 3.5±<br>0.4 | 125.9±<br>9.5 | 210.8±17.7<br>p=<br>0.0007 | 10.3±<br>0.5 | 22.5±1.4<br>p=<br>0.00001 | 4.4±<br>0.7 | 29.9±2.7<br>p=<br>0.00002 | 138.7±9.3 pA<br>(at -120 mV)<br>35.5±2.7<br>(at -50 mV), n=10 |
| **Pitx3 DS**<br>**D1-MSN**<br>**PN18-20**<br>**(1% DA)** | -81.1±0.5<br>n=9 | -78.2±0.5<br>p=<br>0.00001 | 2.9±<br>0.3 | 126.1±<br>4.7 | 195.5±7.9<br>p=<br>0.00001 | 10.6±<br>0.4 | 18.7±1.3<br>p=<br>0.00003 | 4.4±<br>0.6 | 26.4±1.7<br>p=<br>0.00000 | 100.3±4.73 pA<br>(at -120 mV)<br>15.6±1.51 pA<br>at -50 mV, n=9 |
| **Pitx3 NAc**<br>**D1-MSN**<br>**PN18-20**<br>**(50% DA)** | -81.8±0.8<br>n=6 | -81.0±0.8<br>p=<br>0.00112 | 0.8±<br>0.1 | 122.8±<br>5.8 | 137.5±5.9<br>p=0.001 | 10.3±<br>0.6 | 12.1±0.6<br>p=<br>0.00313 | 4.1±<br>1.3 | 9.1±2.1<br>p=<br>0.0054 | ND |
| **WT DS**<br>**D1-MSN**<br>**PN28-30**<br>**(100% DA)** | -81.2 ± 0.7<br>n=6 | -79.9±0.7<br>p=0.0009 | 1.2±<br>0.2 | 104.5±<br>6.8 | 120.7 ± 5.9<br>p=0.0008 | 8.9±<br>0.7 | 11.1 ±0.7<br>p=0.0001 | 2.8±0.<br>7 | 10.1±1.5<br>p=<br>0.003 | ND |
| **TH KO DS**<br>**D1-MSN**<br>**PN28-30**<br>**(0% DA)** | -81.4±0.6<br>n=5 | -78.3±0.4<br>p=<br>0.0018 | 3.1±<br>0.5 | 105.7±<br>4.5 | 179.4±7.6<br>p=0.0004 | 8.7±<br>0.6 | 18.7±0.7<br>p=0.0001 | 3.8±<br>1.4 | 42.6±5.7<br>p=<br>0.0010 | ND |
| **Pitx3 DS**<br>**D1-MSN**<br>**PN28-30**<br>**(1% DA)** | -83.1±1.1<br>n=6 | -80.7±1.1<br>p=<br>0.00022 | 2.6±<br>0.3 | 101.5±<br>5.7 | 192.3±3.5<br>p=<br>0.00004 | 9.9±<br>0.6 | 19.1±1.4<br>p=0.0007 | 2.7±<br>0.6 | 35.0±3.1<br>p=<br>0.00034 | ND |

### 3.2. DA hyperactively increases D1-MSN excitability in the <u>DS</u> in TH KO mice with total DA loss

Next, we examined how DA/D1 agonism affects the intrinsic excitability of D1-MSNs in parkinsonian TH KO mice (**Fig. 1C**) in which DA (derived from injected L-dopa) or D1R agonism produces profound motor stimulation (Kim et al. 2000). We first performed current clamp experiments followed by current clamp experiments; these 2 types of experiments not only provide independent information on neuronal excitability but also verify each other, thus increasing data reliability. Again, to increase recording quality and data reliability, we started with PN18-21 mice that yield more viable brain slices and hence high quality recordings.

#### Current clamp experiments

To examine the potential effects of DA/D1 agonism’s effects on D1-MSN intrinsic excitability, we first made current-clamp recording in D1-MSNs in the dorsal striatum in TH KO mice. To avoid the potential confound of synaptic activity, GABA_A_ blocker picrotoxin and NMDA and non-NMDA receptor blockers AP5 and DNQX were added to the bathing solution in this experiment. To monitor membrane properties at different membrane potentials, determine whole-cell input resistance (R_In_) and evoke spike firing, we injected a series of current pulses (starting at -140 pA and increasing +20 pA per step) into D1-MSNs. Upon hyperpolarizing current injection, these D1-MSNs showed an inward rectification typical of striatal MSNs (**Fig. 3A**) (Nisenbaum and Wilson 1995). Upon injection of strong depolarizing currents, action potentials were evoked in D1-MSNs when the membrane potential reached around –42 mV in TH KO mice (**Fig. 3A**); the action potential peak commonly reached around +30 mV with a duration of about 2.0 ms at the base (recorded at +31 °C) -- these characteristics are typical for MSNs.

**Fig. 3.**
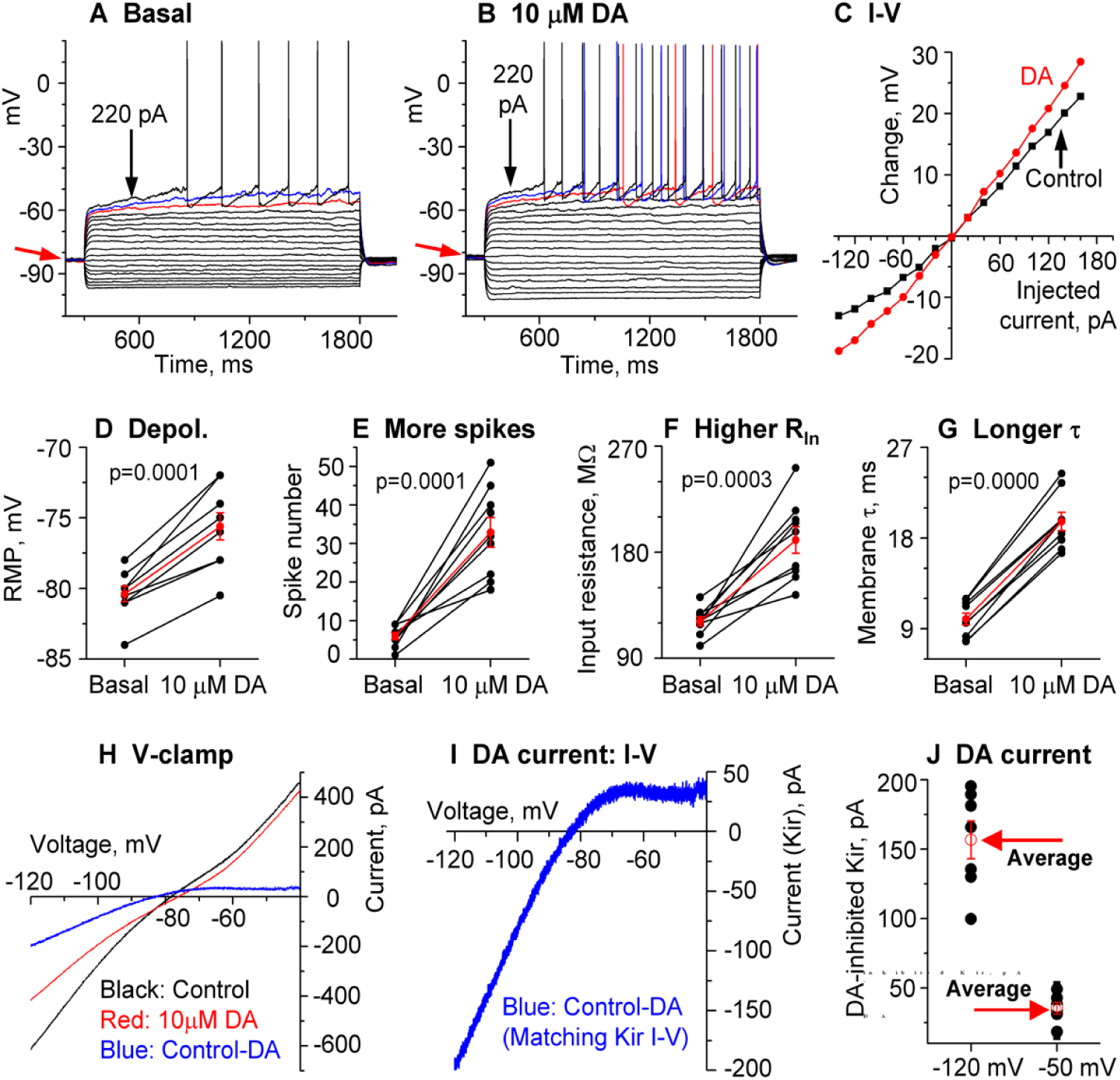
DA hyperactively increases D1-MSN excitability in the **dorsal striatum (DS) in PN18-20 TH KO** mice with total DA loss. **A, B, C.** An example D1-MSN receiving hyperpolarizing and depolarizing current injection before (**A**) and during (**B**) bath application of 10 μM DA and the quantification of membrane potential changes (**C**). **D, E, F, G.** Pooled data showing the DA effect on RMP (**D**), action potential firing (**E**), RIn (**F**), and τ (**G**) in 8 D1-MSNs. P values are from paired t-tests. **H, I.** Example voltage ramp (150 mV/s)-induced K currents (recorded in 1 μM and 0 mM Ca) in an D1-MSN before (control black) and during 10 μM DA (red) (**H**); the difference caused by DA (blue) was extracted by subtraction (**I**) and I–V profile of this DA-sensitive current is typical of Kir. **J.** Pooled data of the DA-sensitive Kir currents at –120 mV and –50 mV in 7 D1-MSNs with the mean ± se plotted in red indicated by the red arrows.

After obtaining a stable baseline recording, we bath-applied 10 μM DA and detected a consistent depolarization in D1-MSNs and this effect was larger in TH KO mice than in WT mice. As shown in **Fig. 3A-C**, and **Table 1**, bath application of 10 μM DA caused a 4.8±0.6 mV depolarization in DS D1-MSNs in TH KO mice, depolarizing them from their RMP at –80.4 ± 0.6 mV to –75.6 ± 1.0 mV under 10 μM DA (p=0.00003, t=–8.63, DoF=8, paired t-test) (**Fig. 3A, B, D**; **Table 1**); simultaneously, bath application of 10 μM DA also increased the whole cell R_In_ from the basal 121.3 ± 3.9 MΩ to 190.4 ± 11.3 MΩ under 10 μM DA (measured by a –20 pA current from the RMP), a 57.2% increase (p=0.0003, t=–6.05, DoF=8, paired t-test) (**Fig. 3F**, **Table 1**); the membrane τ was also increased from the basal 10.0±0.6 ms to 19.6±0.9 ms under 10 μM DA (p=0.0000, t=–17.6, DoF=8, paired t-test) (**Fig. 3G**, **Table 1**) ― this is expected because the membrane τ is proportional to whole-cell R_In_. The increase in the whole-cell R_In_ became larger when the cell membrane potential was more hyperpolarized (**Fig. 3C**), indicative of a potential inhibition of an inwardly rectifying current or Kir (this will be confirmed by voltage clamp experiments described below).

Because of the DA-induced increase in R_In_, DA increased the current injection-induced depolarization and the evoked spikes, but did not change the membrane potential threshold for action potential triggering or waveform. Under the basal condition, ≥180 pA was often needed to evoke spikes (**Fig. 3A**); during 10 μM DA, 140 pA was often sufficient to evoke spikes such that 180 pA evoked many more spikes than under the control condition (**Fig. 3B, E**): 6.0±0.9 spikes were evoked by injected current pulses up to 180 pA under control, whereas 32.9±3.9 spikes were evoked by the same current pulses during 10 μM DA (p=0.0001, t=–7.06, DoF=8, paired t-test) (**Fig. 3A,B,E; Table 1).**

These DA excitatory effects (RMP depolarization, increases in R_In_ and membrane τ in DS D1-MSNs in DD mice were also larger than those in WT mice (evident in the listed numbers in **Table 1** and also supported by ANOVA testing).

#### Voltage clamp experiments

The voltage response profile in our current clamp data shown above suggest that DA may be inhibiting Kir in D1-MSNs (**Fig. 3A-C**). To further establish this conclusion, here we performed voltage clamp experiments to directly examine DA’s potential effects on the Kir in D1-MSNs in TH KO mice. The voltage-dependence or the I-V relationship of the DA-sensitive current was investigated by using a linear voltage ramp ranging from –120 mV to –40 mV. The 10 μM DA-sensitive current was extracted by subtraction. As shown in **Fig. 3H,I**, The 10 μM DA-sensitive current had the characteristics of Kir: it was inwardly rectifying and had a reversal potential near the calculated K reversal potential at –87.5 mV for our extracellular and intracellular solution pair. The amplitude of the DA-sensitive Kir was –156.7±13.5 pA at –120 mV and 36.0±3.7 pA at –50 mV in 7 D1-MSNs in dorsal striatum (**Fig. 3J**, **Table 1**). These characteristics strongly resemble the Kir typically seen in MSNs (Ariano et al. 2005; Shen et al. 2007).

Additionally, DA (10 μM) was without effect on the voltage ramp-evoked currents in the presence of 200 μM BaCl_2_, a known Kir blocker (Hibino et al. 2010; Kubo et al. 2005; Nisenbaum and Wilson 1995; Shen et al. 2007), further confirming that the DA-sensitive current was Kir (**Fig. 4**), and also indicating that Kir was the dominant outward or K current inhibited by D1R activation. It is clear that 200 μM BaCl_2_ produced a total or nearly total inhibition of Kir, thus occluding the partial Kir inhibition normally produced by DA/D1R agonism.

**Fig. 4.**
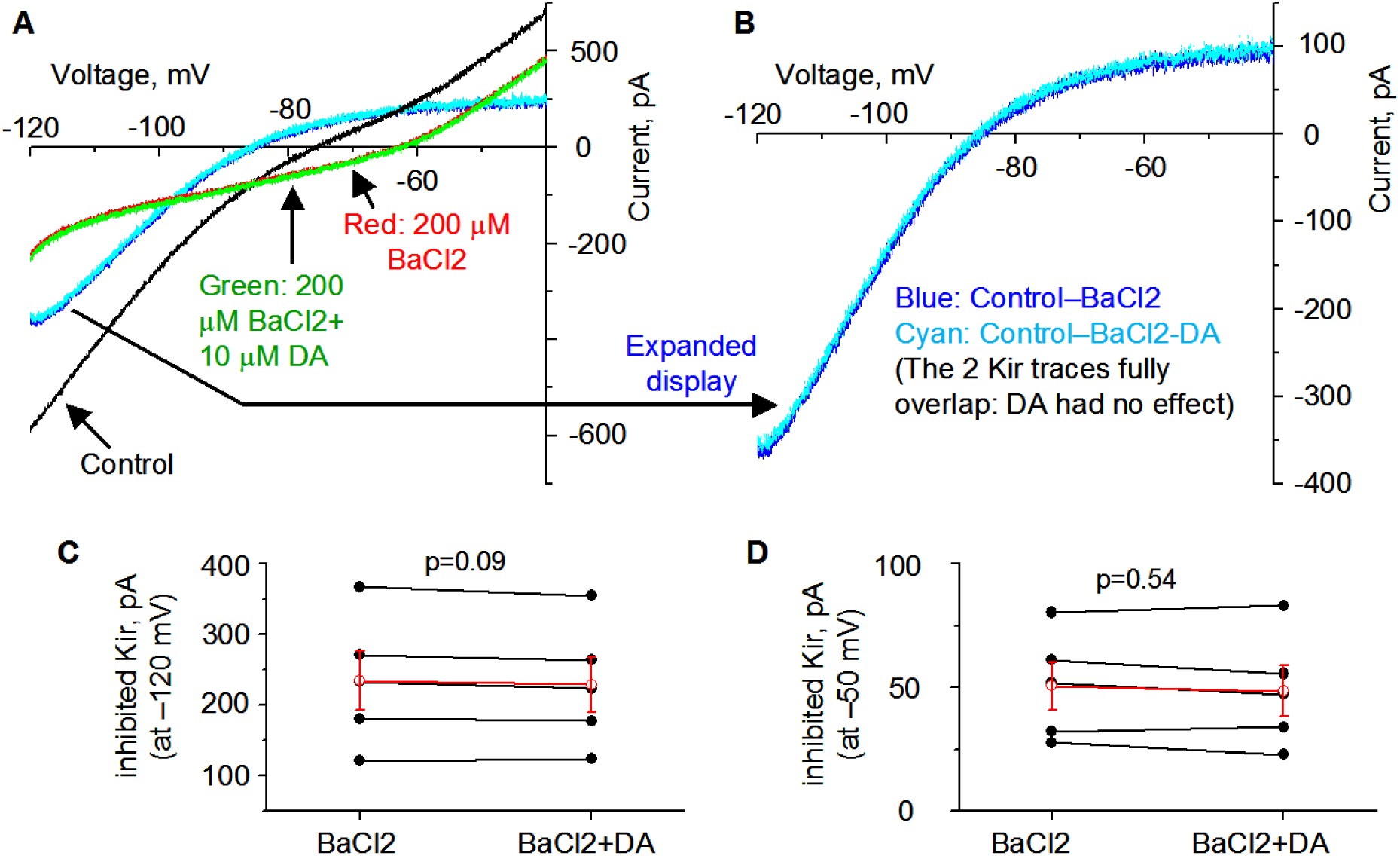
Ba inhibits Kir and occludes DA inhibition of Kir in D1-MSN in DS in **PN18-20 TH KO** mice. **A,B.** Example voltage ramp (150 mV/s)-induced K currents (recorded in 1 μM TTX and 0 mM Ca) in a D1-MSN before (control black) and during 200 μM BaCl2 (red) and during 200 μM BaCl2 with 10 μM DA (green) (**A**); differences caused by BaCl2 (blue) and by BaCl2 with DA (cyan) were extracted by subtraction (**B**) and I–V profiles of this Ba/Ba+DAsensitive current are typical of Kir. **C.** Pooled data of the DA -sensitive Kir currents at –120 mV during 200 μM BaCl2 and during 200 μM BaCl2 with10 μM DA. **D.** Pooled data of the DA -sensitive Kir currents at –50 mV during 200 μM BaCl2 and during 200 μM BaCl2 with10 μM DA. Red symbols and lines are the mean ± se.

### 3.3. DA agonism hyperactively increases D1-MSN excitability in the <u>NAc</u> in TH KO mice with total DA loss

Like the DS, NAc receives intense DA innervation, NAc DA activity has many important functions: motivation, reward, emotion and limbic functions. Thus, understanding the cellular and ion channel mechanisms of DA regulation and dysregulation of NAc neurons is important. Like the DS, the effects DA on MSNs in the NAc were also controversial (Perez et al. 2006; Humphries and Prescott 2010). Here we addressed this long-standing question. We focused on NAc core identified by using the anterior commissure as the landmark.

#### Current clamp experiments

Like D1-MSNs in the DS, upon bath application of 10 μM DA, the RMP was modestly but consistently depolarized by 3.5 ± 0.4 mV, from –81.0 ± 0.5 mV under control condition to –77.5 ± 0.6 mV under 10 μM DA (p=0.00003, t=–9.499, DoF=7, n=8 cells, paired t-test) (**Fig. 5A,B,C**), the evoked spikes numbers were increased from 4.4 ± 0.7 spikes under control to 29.9 ± 2.7 under 10 μM DA (n=8 cells, p=0.00002, t=–10.5045, DoF=7, paired t-test) (**Fig. 5E**, **Table 1**), the whole cell R_In_ was increased from 125.9 ± 9.5 MΩ under basal condition to 210.8 ± 17.7 MΩ under 10 μM DA (n=8 cells, p=0.0007, t=–5.1586, DoF=7, paired t-test, **Fig. 5F**, **Table 1**); and likely as a consequence of increased whole cell R_In_, the membrane charging time constant τ was also increased (10.3±0.5 ms during baseline, 22.5±1.5 ms under 10 μM DA, (n=8 cells, p=0.0007, t=–5.1586, DoF=7, paired t-test) (**Table 1**). These effects were identical to those in DS (compare **Fig. 3** and **Fig. 5**).

**Fig. 5.**
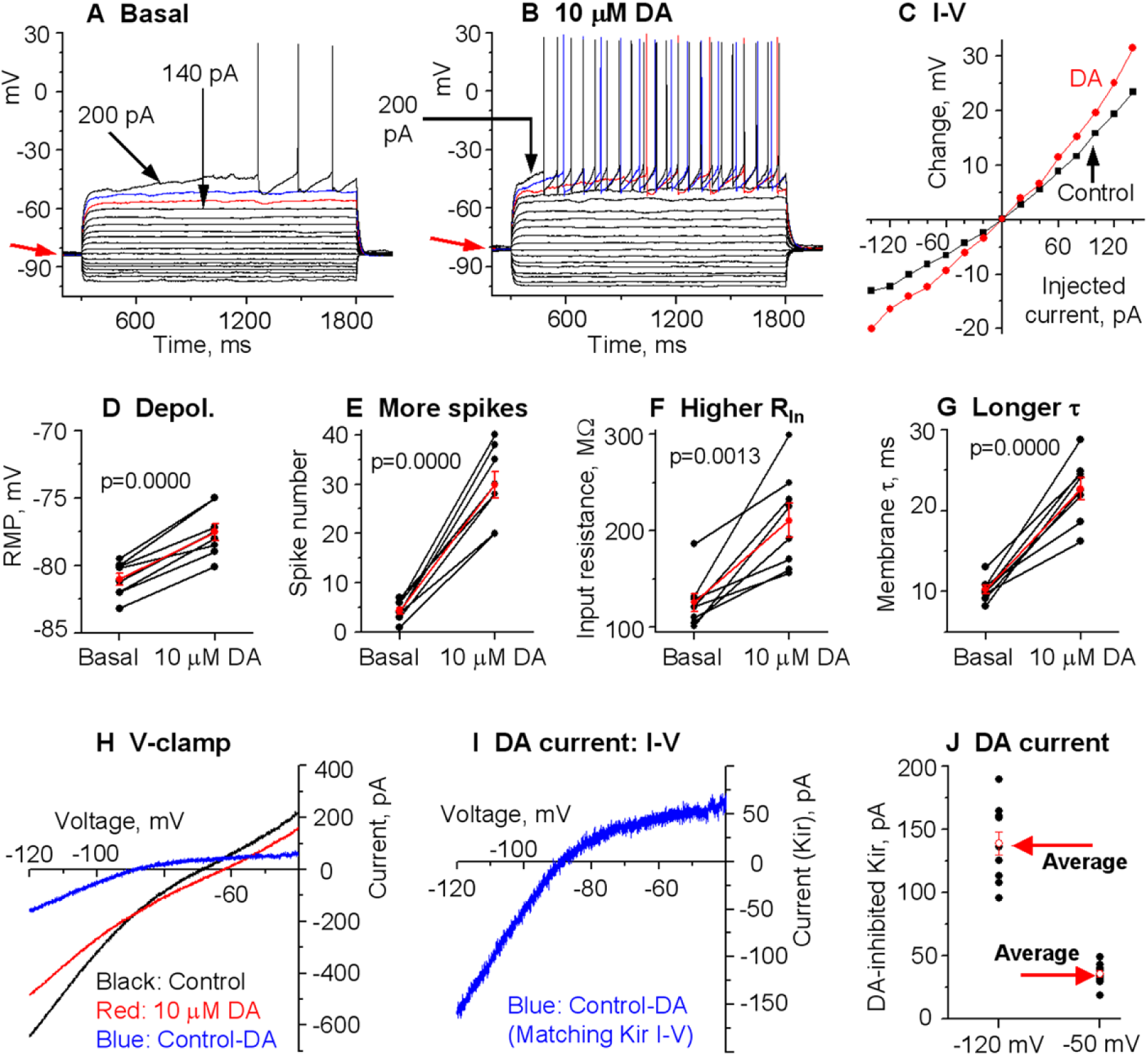
DA hyperactively increases D1-MSN excitability in the nucleus accumbens (**NAc**) in **PN18-20 TH KO** mice with total DA loss. **A, B, C.** An example D1-MSN receiving hyperpolarizing and depolarizing current injection before (**A**) and during (**B**) bath application of 10 μM DA and the quantification of membrane potential changes (**C**). **D, E, F, G.** Pooled data showing the DA effect on RMP (**D**), action potential firing (**E**), RIn (**F**), and τ (**G**) in 8 D1- MSNs. p values are from paired t-tests. **H, I.** Example voltage ramp (150 mV/s)-induced K currents (recorded in 1 μM and 0 mM Ca) in an D1-MSN before (control black) and during 10 μM DA (red) (**H**); the difference caused by DA (blue) was extracted by subtraction (**I**) and I–V profile of this DA-sensitive current is typical of Kir. **J.** Pooled data of the DA-sensitive Kir currents at –120 mV and –50 mV in 10 D1-MSNs with the mean ± se plotted in red indicated by the red arrows.

#### Voltage clamp experiments

Our current clamp data presented above strongly indicate that DA/D1R agonism may be inhibiting Kir in D1-MSNs in NAc in TH KO mice (**Fig. 5A-C**). Here we performed voltage clamp experiments to directly determine DA/D1R agonism’s potential effects on the Kir in NAc D1-MSNs in parkinsonian mice. We evoked Kir by using the same linear voltage ramp from –120 mV to –40 mV. As shown in **Fig. 5H, I**, the 10 μM DA-sensitive/inhibited current, extracted by subtraction, was inwardly rectifying and had a reversal potential near the calculated K reversal potential at –87.5 mV; the amplitude of this DA-sensitive current was – 138.7 ± 9.3 pA at –120 mV and 35.5 ± 2.7 pA at –50 mV in 10 D1-MSNs in NAc (**Fig. 5**; **Table 1**). These results indicate that DA inhibits a current with characteristics of Kir in D1-MSNs in NAc in TH KO mice.

### 3.4. Hyperactive DA responses in D1-MSNs remain in more mature TH KO mice

We next examined DA effects in more mature PN28-30 mice. This is physiologically relevant and also because a recent study reported that DA deficiency increased D1-MSN intrinsic excitability around PN28 and thereafter but not at PN18 (Lieberman et al. 2018). We found that the intrinsic membrane properties of D1-MSNs matured apparently in a similar or identical manner in DA innervation-intact mice and DA-denervated mice such that the baseline intrinsic membrane properties remained similar in DA innervation-intact mice and DA-denervated mice. Specifically, while there were some quantitative changes such as a decrease in whole cell R_In_ due to normal increased Kir expression as the animal becomes more mature, these parameters of the intrinsic membrane properties remained similar in D1-MSNs in DA innervation-intact mice and DA-denervated mice, and the hyperactive DA responses in D1-MSNs in DA-denervated striatum remained (**Fig. 6**; **Table 1**).

**Fig. 6.**
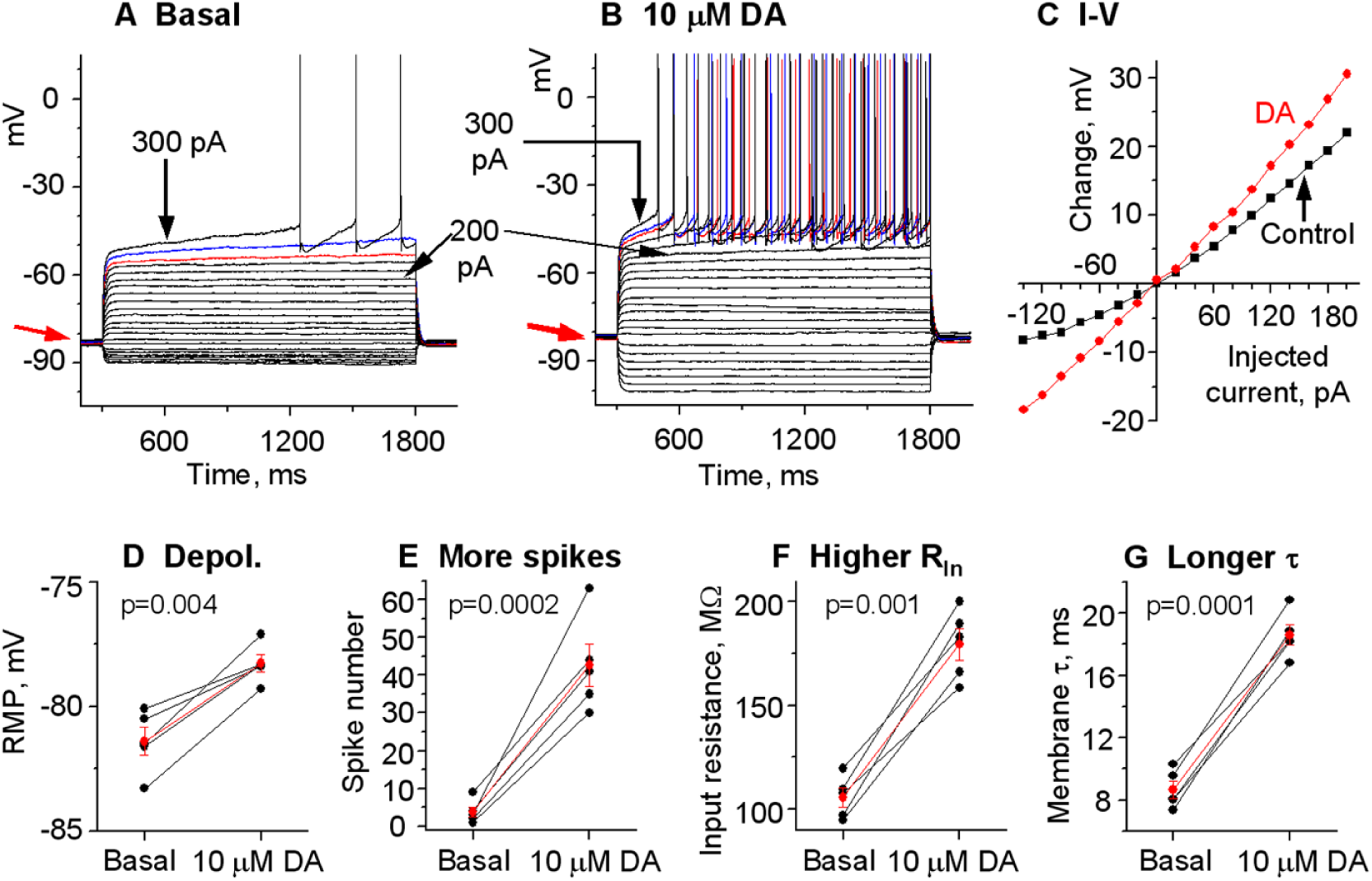
Hyperactive DA responses in D1-MSNs remain in the dorsal striatum in more mature PN28-30 **TH KO** mice. **A, B, C.** An example D1-MSN receiving hyperpolarizing and depolarizing current injection before (**A**) and during (**B**) bath application of 10 μM DA and the quantification of membrane potential changes (**C**). **D, E, F, G.** Pooled data showing the DA effect on RMP (**D**), action potential firing (**E**), RIn (**F**), and τ (**G**) in 5 D1-MSNs. Red symbols and lines are the mean ± se. p values are from paired t-tests.

### 3.5. Gradient hyperactive DA response in D1-MSNs in Pitx3Null mice with gradient DA denervation

#### Hyperactive DA enhancement of D1-MSN excitability in the dorsal striatum (DS) in Pitx3Null mice

Pitx3Null mice have a gradient striatal DA denervation (**Fig. 1A,B**) with the DA loss in the very dorsal DS being ∼ 99% and the NAc DA loss being about 50% [weak to non- supersensitivity, hence no DA receptor functionality and no triggering of c-fos expression upon L-dopa stimulation (Zhong et al. 2023)], resembling the DA loss pattern in PD patients and thus providing an outstanding opportunity for studying potentially gradient DA effects in parkinsonian striatum.

We first used PN18-21 Pitx3Null mice and recorded D1-MSNs in the very dorsal part (within 300 μm from the corpus callosum) of DS where DA denervation is about 99% and D1Rs are hyperfunctional (Kim et al. 2000; Keefe and Gerfen 1996; Paul et al. 1992; Zhong et al. 2023). In current clamp, upon bath application of 10 μM DA, the RMP was depolarized from – 81.1±0.5 mV under control condition to –78.2±0.5 mV under 10 μM DA (n=9 cells, p=0.00001, t=–9.4784, DoF=8, paired t-test) (**Fig. 7A, B and D**, **Table 1**), the evoked action potential numbers were increased from 4.4±0.6 to 26.4±1.7 (n=9 cells, p=0.00000, t=–14.758, DoF=8, paired t-test) (**Fig. 7E**, **Table 1**), the whole cell R_In_ was increased from 126.1±4.7 MΩ under basal condition to 195.5±7.9 MΩ under 10 μM DA (n=9 cells, p=0.00001, t=–9.3708, DoF=8, paired t-test, **Fig. 7F**; **Table 1**), likely as a consequence of increased whole cell R_In_, the membrane charging time constant τ increased from 10.6±0.4 ms during baseline to 18.7±1.4 ms under 10 μM DA (n=9 cells, p=0.00003, t=–8.3786, DoF=8, paired t-test) (**Fig. 7G**; **Table 1**).

**Fig. 7.**
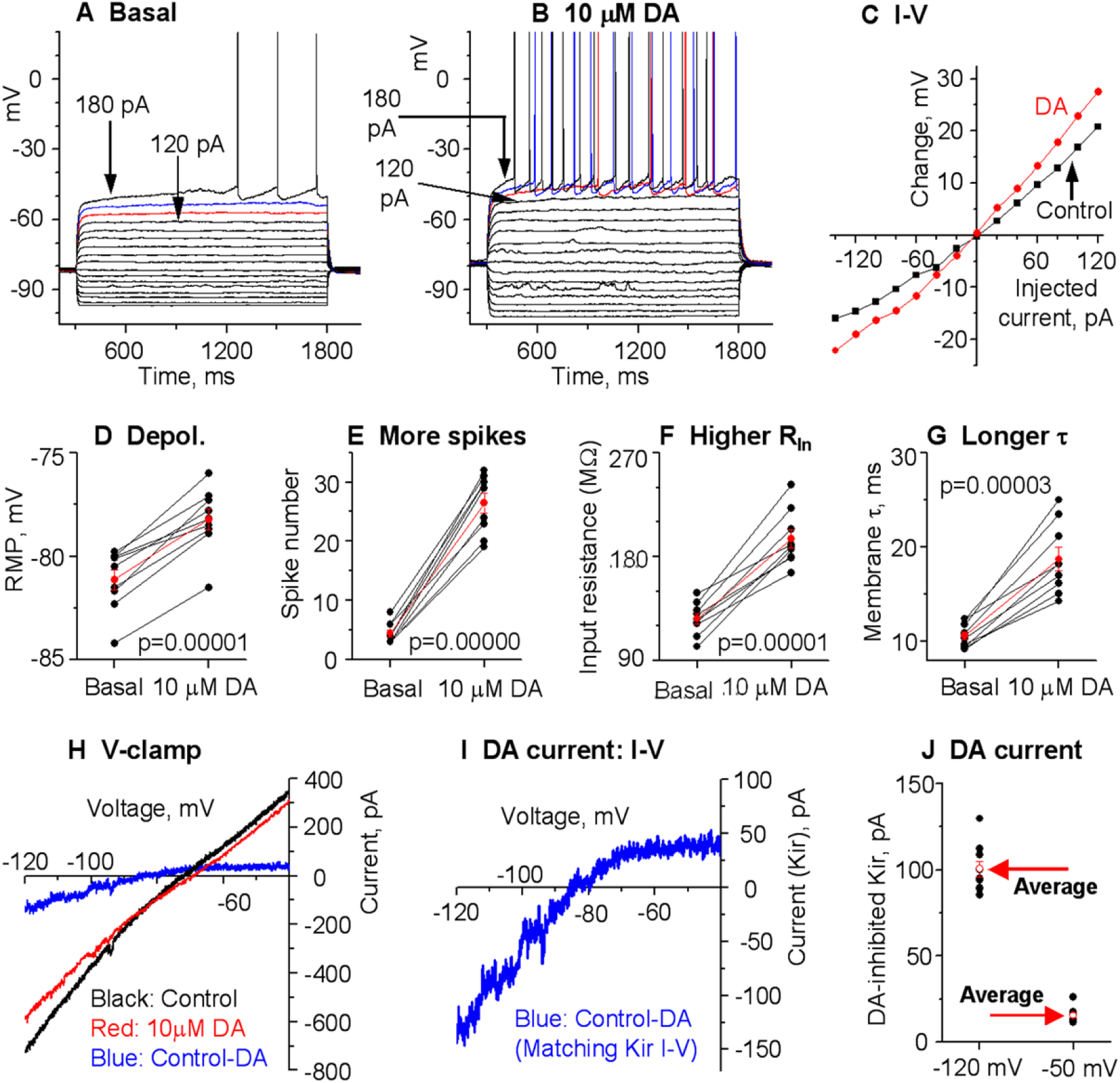
DA hyperactively increases D1-MSN excitability in the **dorsal striatum (DS) in PN18-20 Pitx3Null** mice with severe DA loss in DS. **A, B, C.** An example D1-MSN receiving hyperpolarizing and depolarizing current injection before (**A**) and during (**B**) bath application of 10 μM DA and the quantification of membrane potential changes (**C**). **D, E, F, G.** Pooled data showing the DA effect on RMP (**D**), action potential firing (**E**), RIn (**F**), and τ (**G**) in 9 D1-MSNs. p values are from paired t-tests. **H, I.** Example voltage ramp (150 mV/s)-induced K currents (recorded in 1 μM and 0 mM Ca) in an D1-MSN before (control black) and during 10 μM DA (red) (**H**); the difference caused by DA (blue) was extracted by subtraction (**I**) and I–V profile of this DA-sensitive current is typical of Kir. **J.** Pooled data of the DA-sensitive Kir currents at –120 mV and –50 mV in 9 D1-MSNs with the mean ± se plotted in red indicated by the red arrows.

We further performed voltage clamp experiments to directly determine DA’s potential effects on Kir in D1-MSNs in Pitx3Null mice. To evoke Kir currents, we used the same linear – 120 mV to –40 mV voltage ramp. The 10 μM DA-sensitive current was extracted by subtraction. As shown in **Fig. 7**, the 10 μM DA-inhibited current was inwardly rectifying and had a reversal potential near the calculated K reversal potential at –87.5 mV, matching the characteristics of Kir. The amplitude of the 10 μM DA-inhibited Kir was –100.3±4.7 pA at –120 mV and –15.6±1.5 pA at –50 mV in D1-MSNs in DS in Pitx3Null mice (**Fig. 7**; **Table 1**).

We next examined DA effects in more mature PN28-30 Pitx3Null mice. Data from more mature animals are physiologically important though more difficult to obtain. Additionally, a recent study reported that the DA deficiency in Pitx3Null mice affected the developmental maturation of D1-MSNs such that D1-MSNs in PN28 or older Pitx3Null mice had more depolarized RMP and higher R_In_ than in WT mice, but these intrinsic excitability parameters were normal at PN18 (Lieberman et al. 2018). We found that the basal RMP, R_In_, τ and spike firing in D1-MSNs in PN28-30 Pitx3Null mice were similar to those in PN28-30 WT mice (**Fig. 8A-G**). Furthermore, hyperactive DA responses in DS D1-MSNs remain in more mature Pitx3Null mice: 10 μM DA depolarized D1-MSNs in the DS in PN28-30 Pitx3Null mice, increased their R_In_, τ and evoked spike firing (**Fig. 8A-G)**.

**Fig. 8.**
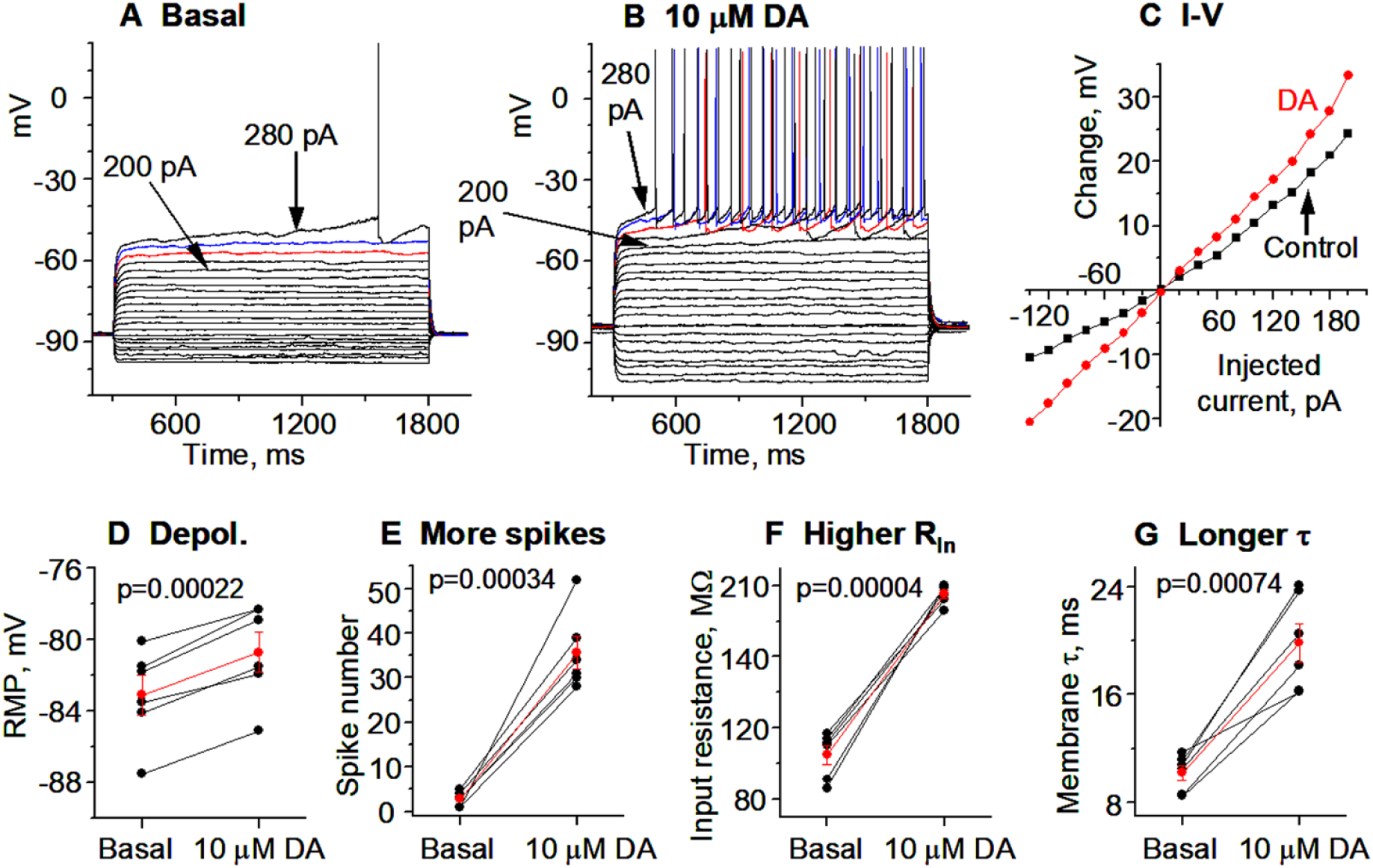
Hyperactive DA responses in D1-MSNs remain the **dorsal striatum** in more mature PN28-30 **Pitx3** mice. **A, B, C.** An example D1-MSN receiving hyperpolarizing and depolarizing current injection before (**A**) and during (**B**) bath application of 10 μM DA and the quantification of membrane potential changes (**C**). **D, E, F, G.** Pooled data showing the DA effect on RMP (**D**), action potential firing (**E**), RIn (**F**), and τ (**G**) in 6 D1-MSNs. Red symbols and lines are the mean ± se. p values are from paired t-tests.

#### Normal DA agonistic enhancement of D1-MSN excitability in the moderately DA- denervated NAc in Pitx3Null mice

In PD, the ventral striatum or nucleus accumbens (NAc) retains significant amounts of DA innervation during middle PD and even in late stage PD (Hornykiewicz 2001). How DA/D1R agonism affects the intrinsic excitability of NAc D1-MSNs with residual DA denervation in parkinsonian striatum was unknown. To address this question, we examined the effects of bath-applied DA on MSN intrinsic excitability of D1-MSNs in NAc in Pitx3Null mice where the residual DA innervation is significant (**Fig. 1A,B**) and resembles the residual NAc DA innervation in middle stage PD. Under a recording condition that was identical to that used for recording DS D1-MSNs, we detected, in current clamp recording mode, that bath application of 10 μM DA only modestly increased the intrinsic excitability of these NAc D1- MSNs in Pitx3Null mice (**Fig. 9**, **Table 1**), just like the modest DA excitatory effect on D1-MSN intrinsic excitability in DA innervation-intact WT mice (**Fig. 1**).

**Fig. 9.**
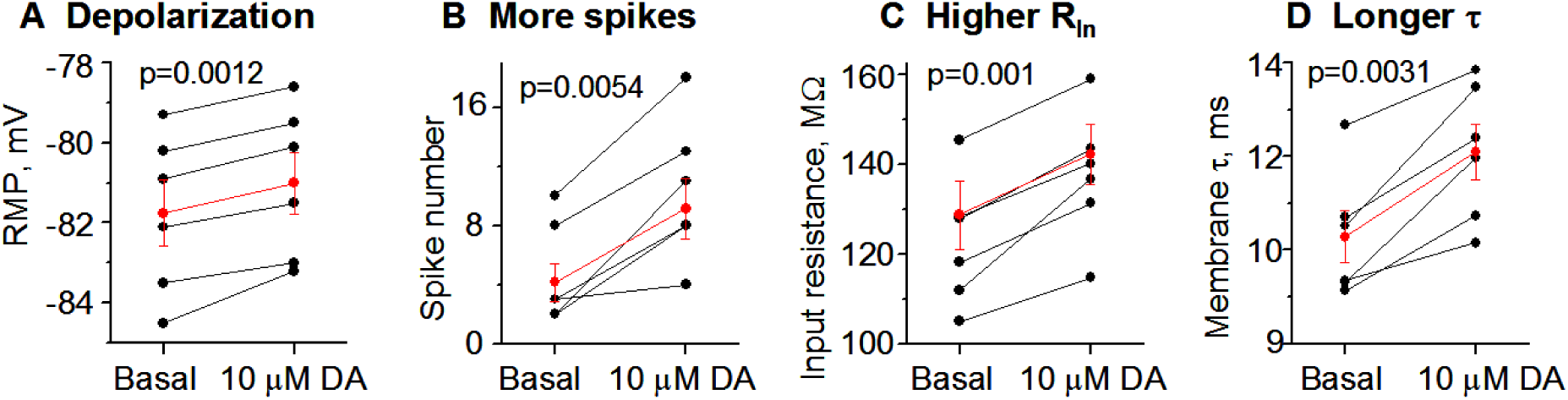
DA normally and modestly increases D1-MSN excitability in the **NAc in PN18-20 Pitx3Null** mice with modest DA loss in NAc. **A, B, C, D.** Pooled data showing the DA–induced depolarization of RMP (**A**), effects on current injection-induced action potential numbers (**B**) input resistance (**C**) and membrane τ (**D**) in 6 D1-MSNs. Red symbols and lines are the mean ± se. p values are from paired t-tests.

Based on the data described in preceding sections and ANOVA testing, it is clear that the baseline membrane properties (RMP, R_In_, τ, and evoked spike firing) of D1-MSNs were similar in the dorsal striatum and NAc in WT, Pitx3Null mice and TH KO mice of mice, indicating that DA loss does not affect the development and maintenance of the membrane properties and the associated ion channels (especially the tonically active Kir, KV channels and NaV channels) of MSNs. In contrast, the DA-induced excitatory effects (depolarization, R_In_ increase, τ increase) were larger in D1-MSNs in the DS and NAc in TH KO mice and the DS in Pitx3Null mice than in DS and NAc in WT mice and NAc in Pitx3Null mice, matching the pattern of DA loss severity, that is, DA’s excitatory effects on D1-MSNs were abnormally high or hyperactive in the striatal region where DA loss was severe.

### 3.6. Local microinjection of BaCl_2_, a Kir inhibitor, into the dorsal striatum, stimulates movement and occludes the motor stimulation of microinjected D1 agonist SKF81297

Data presented above indicate that D1 agonism excites D1-MSNs by inhibiting Kir; literature data indicate that striatal D1 agonist microinjection stimulates movement (Wang and Zhou 2017) and DA receptor-bypassing optogenetic activation of D1-MSNs is known to stimulate movement (Friend and Kravitz 2014; Kravitz et al. 2010; Sippy et al. 2015). Thus, we reasoned that Kir may be a key ion channel mediating D1R agonistic stimulation of motor activity, and hence direct inhibition of Kir in D1-MSNs may stimulate movement and also occludes the motor stimulation of D1R agonism.

To test this idea, we used intra-striatal microinjection of BaCl_2_ and D1 agonist SKF81297. Ba^2+^ is a relatively selective Kir blocker with an IC_50_ around 10 μM (Alagem et al. 2001; Hibino et al. 2010; Kubo et al. 2005), and 100-200 μM BaCl_2_ is often used to completely block Kir in MSNs (e.g. Nisenbaum and Wilson 1995; Shen et al. 2007; Wang and Zhou 2019). We also used unilateral drug injection-induced unilateral rotations as the index of motor stimulation, because (1) rotations can be reliably quantified, and (2) bilateral drug injection often leads to bilaterally spatially asymmetric drug delivery, which interferes with the expression of motor activity and monitoring and quantification. Additionally, we focused on TH KO mice because (1) WT mice have plenty of endogenous DA that prevents the injected agonist from having any major effect, (2) with a total lack of endogenous DA, TH KO mice are likely to produce the strongest and hence most reliable responses to D1R agonism.

As expected, unilateral intrastriatal microinjection of the D1 agonist SKF81297 (0.2 μg in 0.2 μL) induced robust motor stimulation, in the form of contralateral rotations, in TH KO mice, reaching the peak effect at 30 min after injection, stayed at the peak level (∼ 30 rotations/10 min) for the next 30 min – the duration of our monitoring (**Fig. 10, Supplemental video 1**), consistent with our prior studies in a different mouse model of DA deficiency (Wang and Zhou 2017).

**Fig. 10.**
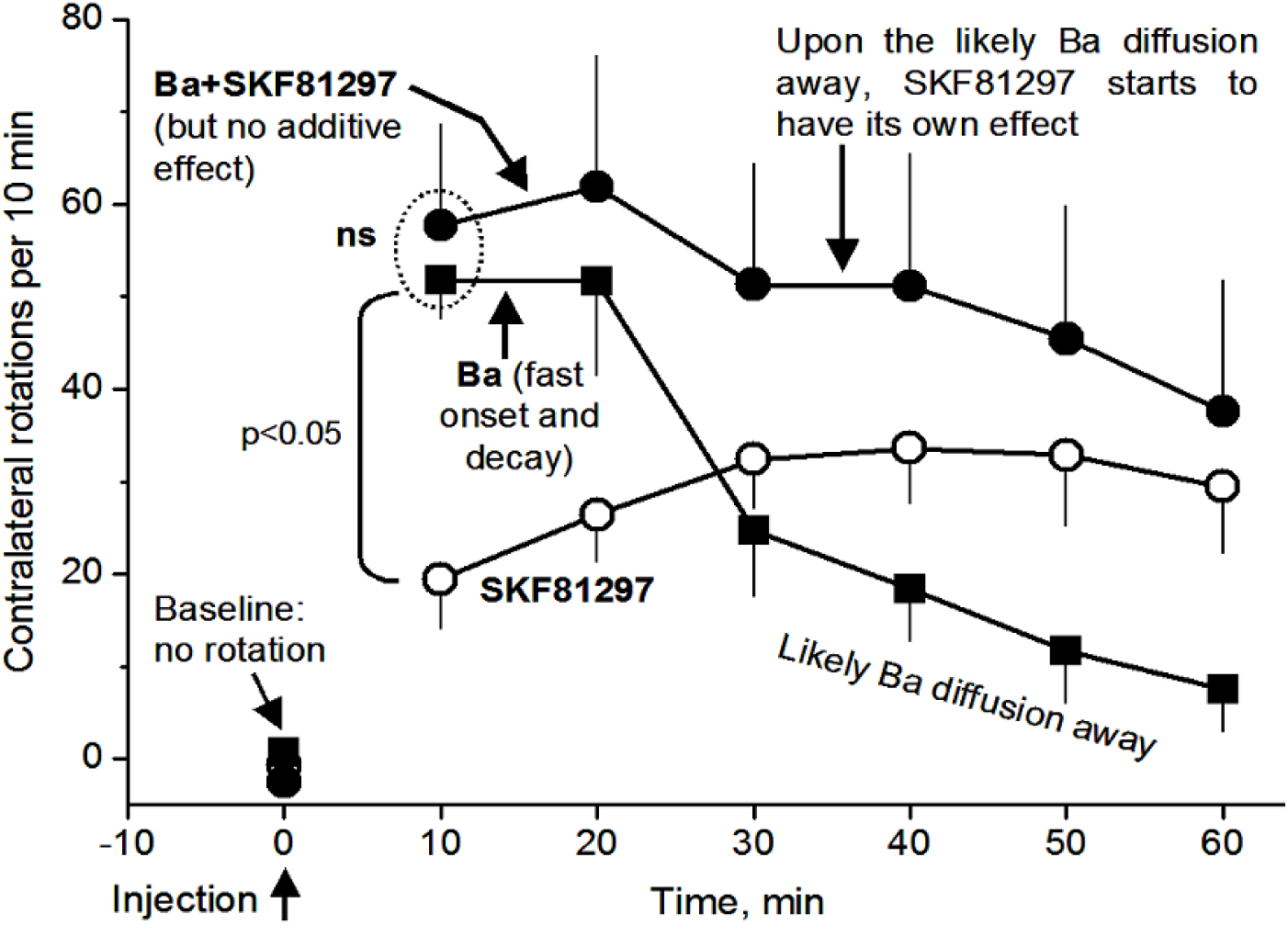
Inhibition of Kir occludes the motor stimulation of D1 agonism. SKF81297 (0.2 μg in 0.2 μL saline) unilateral microinjection into the DS induced contralateral rotations in 5 TH KO mice, see supplemental video 1. BaCl2 (0.2 μg in 0.2 μL) unilateral microinjection into the DS induced robust contralateral rotations in 6 TH KO mice, see supplemental video 2. This Ba effect had a fast onset and also decayed quickly, likely because Ba can diffuse away quickly. Concurrent microinjection of BaCl2 (0.2 μg) and SKF81297 (0.2 μg in the same 0.2 μL together with BaCl2 in 7 TH KO mice triggered only 58 rotations, similar to the 52 rotations triggered by BaCl2 microinjection alone. The data points at 10 min after injection are most pertinent to our question on Kir and motor stimulation and were analyzed with one way ANOVA; data points at and after 30 min after drug injection likely involve differential diffusion of Ba and SKF81297 besides their respective pharmacological effects, and thus were not analyzed statistically.

Furthermore, intrastriatal BaCl_2_ microinjection (0.2 μg in 0.2 μL), in a separate experiment, also induced contralateral rotations that was qualitatively similar to SKF81297- induced contralateral rotations (**Fig. 10, Supplemental video 2**); the Ba effect also occurred faster, reaching the peak effect (∼ 50 rotations/10 min) at 10-20 min after Ba injection, and decayed faster: the decay started at 20 min after Ba injection (**Fig. 10**). The faster rise and decay are consistent with the fact that Ba is a simple divalent cation and can diffuse more easily, whereas SKF81297 is a much more complex molecule and likely diffuses and reaches the maximal sphere of influence more slowly than Ba.

Equally important, as shown in **Fig. 10**, after concurrent microinjection of BaCl_2_ (0.2 μg) and SKF81297 (0.2 μg in the same 0.2 μL together with BaCl_2_) did not produce an additive rotation-stimulating effect; in fact, during 10 min after microinjection, only 58 rotations were triggered, similar to the 52 rotations triggered by BaCl_2_ microinjection alone. This is not because the rotational motor capacity was saturated; these mice can make > 100 rotations/10 min when stimulated strongly. Instead, these results indicate that Ba was blocking and then occluding the same target (i.e. Kir in D1-MSNs) that D1 agonism-cAMP also tried to inhibit. Further, because Ba excites (via inhibiting Kir) both D1-MSNs (**Fig. 4**) and D2-MSNs (Shen et al. 2007), and D2- MSN excitation is motor-inhibitory and counter the motor-stimulating effect of D1-MSN excitation, thus, Ba’s motor-stimulating effect via Kir inhibition in D1-MSNs would be even larger than what we observed here if the counter-effect of D2-MSNs were not there.

## 4. Discussion

Our main findings are that (1) DA activation of D1Rs in D1-MSNs inhibits Kir, leading to substantially increased D1-MSN intrinsic excitability and spike firing, and this DA inhibition of Kir was occluded by BaCl_2_, (2) local BaCl_2_ microinjection-induced likely blockade of Kir stimulated motor activity and also occluded the motor stimulation of D1R agonism, indicating that Kir is a key mediator of DA’s profound behavioral effects, and (3) DA loss does not affect the baseline intrinsic membrane properties but renders D1R agonistic enhancement of D1-MSN intrinsic excitability hyperactive. Our results not only resolve the important and long-standing questions about how DA affects the intrinsic excitability and spike output of D1-MSNs, but also establish that inhibition of Kir in D1-MSNs is the key ion channel mechanism underlying D1R agonism’s profound motor stimulation.

### 4.1. Hyperactive D1R activation inhibits Kir in D1-MSNs and increases their intrinsic excitability in parkinsonian animals displaying hyperactive D1R agonistic motor response

Despite the intense expression of D1Rs and Kir in D1-MSNs, a D1R regulation of Kir and spike firing in D1-MSNs that can be logically interpreted in the context of D1-MSN function in the BG circuit and behavior had not been established until our present study. Employing the strategy of using DA-lacking TH KO mice that likely have larger cellular DA responses than normal mice, we collected robust data in DA-lacking mice that show unambiguously that the activation of cAMP-producing D1Rs inhibits Kir and increases the R_In_ in D1-MSNs in both the DS and NAc, thus substantially increasing the intrinsic excitability of these MSNs. The large amplitude and consistency of the DA excitation detected in both current clamp and voltage clamp strongly indicate that our results are reliable and correct, and also make physiological sense as discussed below. In normal mice with intact striatal DA innervation in the striatum, DA induced excitatory effects in D1-MSNs qualitatively similar to those in parkinsonian mice, although these normal dopaminergic excitatory effects are much smaller.

Our present patch clamping results from in vitro brain slices are consistent with our published in vivo spike recording and in vivo DA drug microinjection studies indicating that D1 agonism in the DA-denervated dorsal striatum triggers hyperactive spike firing in D1-MSNs and hyperactive motor activity (Sagot et al. 2018; Wang and Zhou 2017). Our present results are also consistent with Parker et al. (2018) reporting that Ca indicator-based multi-cell activity recording indicates that D1-MSNs responded hyperactively to L-dopa in 6-OHDA DA denervation PD mice, and with Ryan et al. (2018) reporting that in vivo extracellular spike recording indicated that L-dopa overexcited D1-MSNs in 6-OHDA DA denervation PD mice. Thus, it is reasonable to conclude that DA/D1 agonism may directly increase D1-MSN spiking activity that in turn stimulates motor activity, although network effects may also contribute to the overall spiking activity following DA/D1 agonism administration. Our present data on DA effects on Kir obtained in brain slices are consistent with recent spike data recorded from awake parkinsonian animals that show that L-dopa or D1 agonism stimulated D1-MSN firing hyperactively in DA-depleted striatum (Singh et al. 2015, 2016; Parker et al. 2018; Ryan et al. 2018; Sagot et al. 2018).

Our finding that forskolin mimicked the Kir-inhibition effect of DA in D1-MSNs is consistent with the canonical D1R signaling mechanism: The main D1R signaling pathway, in both normal and parkinsonian striatum, is to increase cAMP and activate the cAMP-PKA signaling pathway in D1-MSNs (Beaulieu et al. 2011; Corvol et al. 2001; Herve 2011; Lee et al. 2002; Missale et al. 1998; Stoof and Kebabian 1981; Nagai et al. 2016). Perhaps more importantly, our results that D1R agonism and the likely consequent increase of intracellular cAMP inhibit Kir in D1-MSNs match our recent study that cAMP-producing Gs-DREADD activation inhibits Kir and increases intrinsic excitability in MSNs (Wang and Zhou 2019). Additionally, our results are consistent with D1R’s inhibition of Kir in cortical pyramidal neurons (Dong et al. 2004).

Another line of evidence supporting the correctness of our data is the internal consistence and control in our data collected from D1-MSNs from WT mice, Pitx3Null mice and TH KO mice are consistent with and support each other, for example, due to the substantial residual DA innervation in the NAc in Pitx3Null mice, DA response is expected to be normal and similar to those in WT but different from those in DS in Pitx3Null and those in DS and NAc in TH KO mice. Our data show that this is indeed the case.

Prior studies examined DA regulation of D1-MSN intrinsic excitability in normal DA innervation-intact striatum, but the results were confounded. In WT mice, Planert et al. (2013) detected a 60 μM DA-induced 30 pA inward current (the ion channel was not identified) in identified DS D1-MSNs but saw no change in input resistance; this result is difficult to interpret because at a fixed membrane potential, an inward current (or any current) requires a change in resistance, according to Ohm’s law. Planert et al. (2013) further reported that 60 μM DA decreased the threshold current to trigger action potentials but increased the AP threshold potential (2 mV more depolarized); these opposing effects are difficult to interpret and potentially non-biological (i.e. they were potentially experimental artifacts): why would DA induce opposing effects on the same D1-MSNs? Podda et al. (2010) reported that in normal NAc in mice, D1 agonism induced an inward current by blocking Kir, but the results were also confounded by the fact that the recordings were made in mixed, unidentified MSNs; consequently, in theory, 50% of the recorded cells should not respond to D1 agonist. Podda et al. (2010) also did not examine DA effects on MSN spike firing. A more recent study (Zhao et al. 2016) reported that D1R agonism increased Kir in identified D1-MSNs and inhibited D1-MSN spike firing; but these results are not compatible with the overall function of D1Rs and the fact that D1-MSN activity stimulates movement (Friend and Kravitz 2014) and D1 agonism in the striatum stimulates movements (Wang and Zhou 2017). Thus, the literature data in this field are conflicting, and thus our present study is a major advance.

We need to note here that although our data clearly show that D1R agonism inhibits Kir in MSNs, D1R agonism may also trigger the upregulation of inward currents that become active at -60 mV or more positive such as the persistent Na current that has been reported to be enhanced by D1R agonism in cortical neurons (Gorelova and Seamans 2015).

### 4.2. DA response intensity in D1-MSNs is dependent on DA denervation severity

We found that in Pitx3Null mice, while DA had hyperactive excitatory effects on D1-MSNs in the DS where DA denervation is very severe, the DA excitatory effects on the intrinsic excitability of D1-MSNs was modest and normal in the NAc where DA denervation is moderate and D1Rs are normal as verified by a lack of L-dopa-induced c-fos expression (Zhong et al. 2023). This finding has 2 lines of importance. The first is that in PD, the DA denervation also has a dorsal-ventral gradient with the DA loss being the severest in the DS and less severe in VS throughout the stages of PD, especially during early and middle stages (Kish et al. 1988; Hornykiewiciz 2011; Kordower et al. 2013). Our present neurophysiological data plus the c-fos data indicate that D1Rs in the central striatum (CS) and VS are probably normal in early and middle stage PD, which may be a factor that L-dopa treatment induces few behavioral problems in these patients. In late stage PD when DA loss becomes severe even in VS, then D1Rs in CS and VS also become abnormal and hyperfunctional, contributing to the dysregulation of brain functions such as cognition, emotion, and limbic functions. The second importance of our Pitx3 NAc data is that they provide an internal control (different DA loss severity leads to different D1R agonistic responses in a logical manner) and indicate the reliability of our data. This is important because of conflicting, probably incorrect data on this topic in the literature.

### 4.3. D1-MSN basal intrinsic membrane excitability is normal in parkinsonian striatum

A potential effect of DA loss on the intrinsic membrane properties such as RMP and R_In_ had been an unsettled important fundamental question with conflicting reports even for the same cell type in the same animal model. In brain slices from the Pitx3Null mice, Lieberman et al. (2018) reported that D1-MSNs were hyperexcitable with a higher R_In_ and more depolarized RMP than in WT mice, but Suarez et al. (2018) reported that, also in brain slices from Pitx3Null mice, D1- MSNs were hyperexcitable (but with a normal R_In_ and RMP) than in WT mice. Additionally, Suarez et al. (2018) reported no change in D1-MSN R_In_ but reported a loss of dendrites and dendritic spines in Pitx3 mice; interpretation of this result is difficult because membrane loss due to dendrite loss should increase R_In_.

In our present study, we found that the basal intrinsic membrane properties and spike firing of D1-MSNs were similar in brain slices from WT mice, TH KO mice and Pitx3Null mice (synaptic receptors were blocked, and DA release in WT mouse brain slices was minimal and dissipated quickly; so we were comparing pure intrinsic membrane excitability). Our present data contradict prior studies reporting increased D1-MSN excitability in 6-OHDA DA-depleted and Pitx3Null parkinsonian striatum in isolated brain slices (Fieblinger et al. 2014; Lieberman et al. 2018; Suarez et al. 2018). These prior studies have a fundamental problem: an increased D1-MSN intrinsic excitability and spiking activity in parkinsonian striatum would increase motor activity (this chain of events is well established, see, for example, Friend and Kravitz 2014; Sippy et al. 2015), contradicting the fact that DA loss leads to akinesia or PD. Additionally, the displayed data in these reports manifest confounding technical issues that might have contributed to the difficult-to-interpret data and conclusions. In contrast, our data are fully consistent with the function of D1-MSNs in the normal striatum and the dysfunction of parkinsonian striatum: In normal striatum, the moderate D1R agonistic enhancement of D1- MSNs facilitates normal motor activity; loss of this D1R facilitation contributes to the reduced motor function in PD; in parkinsonian striatum, D1Rs are hyperfunctional, consequently, the D1R enhancement of D1-MSN excitability and hence D1R facilitation of motor activity are also hyperfunctional, leading to motor function restoration and even motor hyperactivity depending on the dose of D1R agonism. Of course, D2Rs and D2-MSNs also contribute to DA facilitation of motor function, but our focus here is D1Rs and D1-MSNs.

In contrast to the confounded literature data discussed above, our measured basal RMP and R_In_ of D1-MSNs from WT mice, Pitx3Null mice and TH KO mice are consistent with and support each other. Our results are also consistent with in vivo intracellular recordings in MSNs showing that 6-OHDA-induced DA denervation did not alter the intrinsic membrane properties (Pang et al. 2001; Tseng et al. 2001). Further, these neurophysiological data are corroborated by our anatomical data that MSN soma size, dendrite length and dendritic spine number are similar in WT, Pitx3 and TH KO mice (Zhong et al. 2023); thus the two factors that determine the MSN basal R_In_ are similar in normal and parkinsonian striatum: the anatomical structure (hence membrane area) and the basal ion channel activity in the membrane.

Additionally, a new study (Lahiri and Bevan 2020) reported that a 5-pulse (20 Hz, mimicking burst firing) optogenetic stimulation (evoking DA release) increased D1-MSN excitability and this effect ***lasted*** for "at least 10 min," by inhibiting Kv and KCa currents. But it’s firmly established that in normal striatum, the DA axon terminal released DA and its downstream signal last for ∼ 1 s (Rice et al. 2011, Marcott et al. 2018); it is also known that the PKA-sensor-detected DA signal lasting several minutes was due to the slow kinetics of the sensor (Ma et al. 2018; Yapo et al. 2017). Thus, the true nature of this optogenetically induced long-lasting D1-MSN excitation is unknown.

### 4.4. Kir is a key ion channel mediating DA/D1R agonism’s profound motor stimulation

In this study, we found that microinjection of the Kir blocker Ba into the DS triggers contralateral rotation and occludes the motor stimulation of D1R agonism, indicating that Kir is the common target for D1R agonism-induced intracellular signaling in D1-MSNs and direct Ba inhibition and that inhibition of Kir in D1-MSNs can stimulate movement. To our knowledge, this is the first identified, DA-regulated ion channel in BG or D1-MSNs that is directly connected to motor stimulation. Other ion channels such as sodium Nav channels are certainly required for BG/brain to operate generally but not for DA to selectively and specifically regulated that produces motor effects, at least not as established as we have done for Kir in D1-MSNs. Thus, our present study is a groundbreaking study and reveals fundamental neurobiological mechanisms about Kir, its inhibition by D1R activation, and its direct behavioral function.

We also need to discussion this question: since Ba inhibits Kir in both D1-MSNs and D2- MSNs non-selectively, how can Ba stimulate movement? We believe the following is the underlying mechanism. Although Ba inhibits Kir non-selectively, the behavioral consequences of Kir inhibition in D1-MSNs and D2-MSNs should be motor-promoting and motor-inhibiting, respectively, based on the results of optogenetic activation of D1-MSNs and D2-MSNs in the literature (Friend and Kravitz 2014; Kravitz et al. 2010; Sippy et al. 2015). Thus, Ba-induced Kir inhibition in D1-MSNs in a unilateral striatum should stimulate motor function of the ipsilateral striatum and produce contralateral rotation; at the same time, Ba-induced Kir inhibition in D2- MSNs inhibits contralateral rotation. The fact that unilateral Ba injection into the striatum triggered contralateral rotation indicates clearly that Ba inhibition of Kir in D1-MSNs stimulates motor function, and this effect dominates; without the counter-effect in D2-MSNs, Ba’s stimulation of contralateral rotation, via Kir inhibition in D1-MSNs, probably would be larger than we observed here. Our data also indicate that the behavioral inhibitory effects of Ba excitation of D2-MSNs is overcome by the behavioral stimulating effects of Ba excitation of D1-MSNs. This logical interpretation is further supported by our unpublished data (Yuhan Wang and Fu-Ming Zhou) that unilateral microinjection of GABA_A_ receptor blocker bicuculline induced contralateral rotation despite bicuculline’s non-selective blockade of GABA_A_ receptors on all neurons and hence general excitation of all striatal neurons at and near the bicuculline injection site, also indicating that D1-MSN excitation-induced motor stimulation (here in the form of contralateral rotation) can overcome ***D2-MSN excitation***-induced motor ***inhibition***.

Microinjected Ba2+ can also inhibit Kir2 in striatal cholinergic interneurons (Kir2 expression is low in these neurons) and striatal GABA interneurons (Kir2 expression is also relatively low -- that is a key reason MSNs have the most negative membrane potential). But motoric contribution from these interneurons is far smaller than the contributions from D1-MSNs and D2-MSNs, e.g., mice with their striatal cholinergic interneurons ablated can survive and have normal locomotor activity (with some abnormal behaviors) (Kaneko S et al. 2000), and partial ablation of striatal GABA interneurons has only some very modest behavioral effects (Xu et al. 2016); in contrast, MSN slow degeneration in Huntington’s disease causes profound motor and behavioral symptoms and eventually death of the patient (Walker 2007). These vast differences in the functional power of striatal MSNs vs. striatal interneurons are because MSNs comprise at least 90% of striatal neuronal population and are the output neurons of the striatum, whereas striatal interneurons comprise less than 10% of striatal neuronal population, are local interneurons, and exert their function by affecting MSNs. Therefore, striatal MSNs have the primary functional importance in the striatum, whereas striatal interneurons have a secondary functional importance. Thus, Ba effects on striatal interneurons (likely excitation of these neurons) are not likely to be the mediator of Ba-induced contralateral rotation, although a minor contribution is likely (the key mediator is Ba excitation of D1-MSNs, as discussed in the last paragraph).

Our microinjected BaCl_2_ ( 4 mM, 0.2 μL) was unlikely to have interfered with Ca-dependent processes such as glutamate release because, as indicated by the rapid onset and offset of the Ba stimulation of contralateral rotation, injected Ba likely diffused and diluted quickly such that it cannot significantly impede Ca influx. If Ba predominantly inhibits Ca influx and hence glutamate release, then D1-MSNs should be inhibited and there should be no contralateral rotation, or ipsilateral rotation may be triggered, opposite to our observed contralateral rotation.

## 5. Conclusions

In summary, our present study resolves two important questions. First, which is the most critical ion channel D1Rs regulates (inhibit or enhance) in D1-MSNs? Second, does this D1R regulation directly affect behavior? Now we have provided evidence showing that in D1-MSNs, DA activation of D1Rs in D1-MSNs inhibits the highly expressed and tonically active Kir and increases input resistance, intrinsic excitability and spike firing, thus delineating an essential ion channel and cellular mechanism for the intense DA innervation and the heavily expressed D1Rs to regulate the striatum and promote behavior; in Parkinson’s disease, this mechanism is enhanced because of the hyperfunctional D1Rs (although this mechanism is off when DA treatment is off), enabling DA and D1R agonism to profoundly stimulate behavior upon DA treatment (D2Rs and D2-MSNs also participate critically in DA promotion of behavior, see, for example, Wang and Zhou 2017).

## Acknowledgements

This work was supported by NIH/NINDS grant R01NS097671 to FMZ, 1R01-AG058467 to FFL and R01NS120327 to FFL and FMZ.

## Declaration of Interests

The authors declare no competing financial interests.

## Abbreviations used

AP5: d,l-2-amino-5-phosphonovalerate
AC: anterior commissure
cAMP: cyclic adenosine monophosphate
DA: dopamine
D1R: dopamine D1 type receptor
D2R: dopamine D2 type receptor
DNQX: 6,7-dinitro-quinoxaline-2,3-dione
Gs-DREADD: Gs protein-coupled designer receptor exclusively activated by designer drug
DS: dorsal striatum
GABA: gamma amino butyric acid
G_s_: G_αS_ protein
G_olf_: G_αolf_ protein
MSN: medium spiny neuron
dMSN: direct pathway medium spiny neuron
D1-MSN: dopamine D1 receptor-expressing MSN, same as dMSN
iMSN: indirect pathway medium spiny neuron
D2-MSN: dopamine D2 receptor-expressing MSN, same as iMSN
Kir: inwardly rectifying potassium current
KO: knockout
MΩ: Megaohm (10^6^ ohm)
NAc: nucleus accumbens
NMDA: n-methyl-d-aspartate
pA: picoampere (10^-12^ ampere)
PCR: polymerase chain reaction
PD: Parkinson’s disease
R_In_: whole cell input resistance
RMP: Resting membrane potential
τ: membrane time constant
TH: tyrosine hydroxylase
TTX: tetrodotoxin
WT: wild type

## Supplemental video data

**Video 1.**
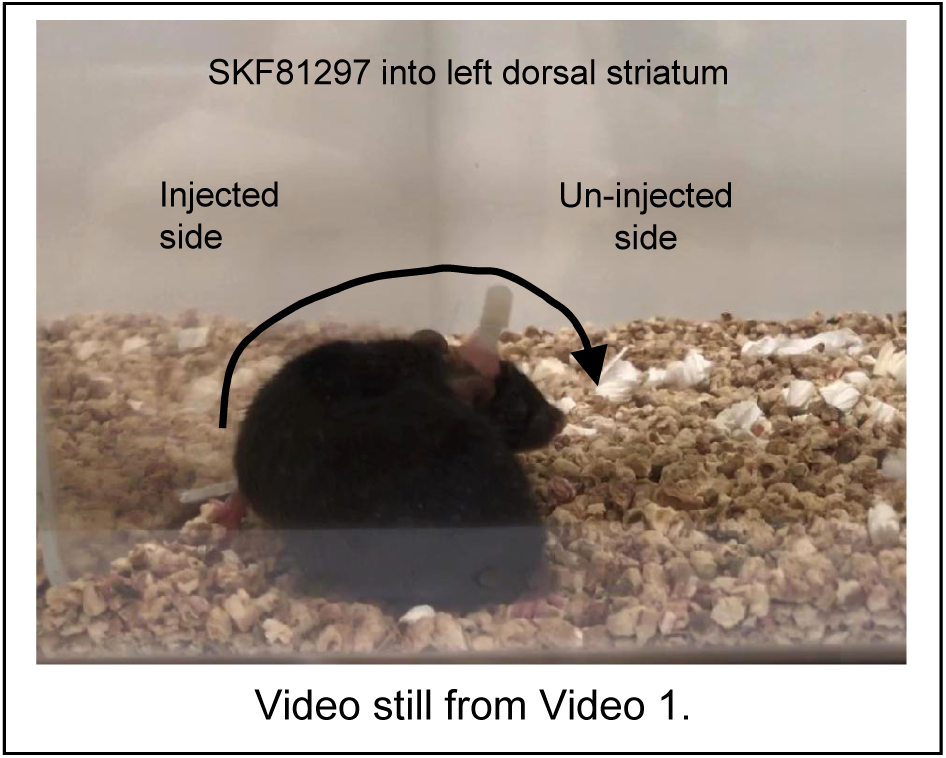
SKF81297 (0.2 μg in 0.2 μL saline) microinjection into the dorsal striatum induced contralateral rotations. This video clip was about <u>25 min after the start of injection</u>. The microinjection tubing had been removed after the 5-min slow microinjection to avoid tissue damage and another 5 min for the injector to remain in place to avoid the reflux of the injected solution.

**Video 2.**
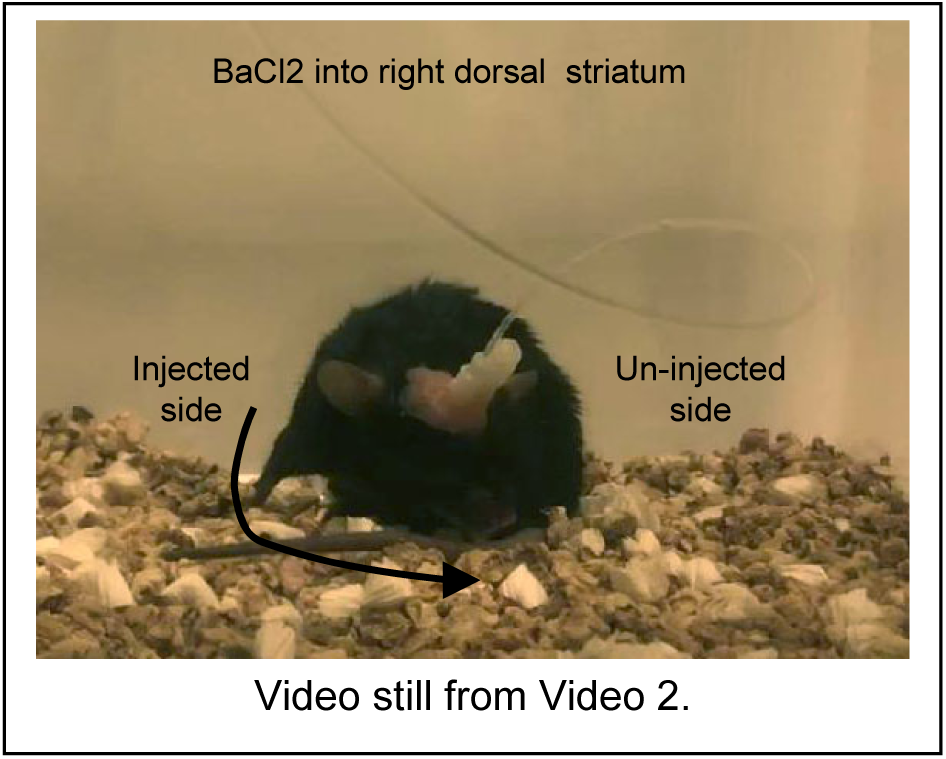
BaCl2 (0.2 μg in 0.2 μL saline) microinjection into the dorsal striatum induced contralateral rotations. This video clip was about <u>8 min after the start of injection</u>. (the onset of BaCl2 effect was faster than that of SKF81297, see also Fig. 10). The microinjection tubing was still connected and visible because the microinjection cannula was allowed to remain 5 min after the 5-min slow microinjection to avoid tissue damage and the reflux of the injected solution.

## Notes

### Competing Interest Statement

The authors have declared no competing interest.

